# Cell type-specific attenuation of brassinosteroid signalling drives stomatal asymmetric cell division

**DOI:** 10.1101/2022.11.26.518021

**Authors:** Eun-Ji Kim, Cheng Zhang, Boyu Guo, Thomas Eekhout, Anaxi Houbaert, Jos R. Wendrich, Niels Vandamme, Manish Tiwari, Claire Simon--Vezo, Isabelle Vanhoutte, Yvan Saeys, Kun Wang, Yuxian Zhu, Bert De Rybel, Eugenia Russinova

## Abstract

In *Arabidopsis thaliana*, the negative brassinosteroid (BR) signalling regulator, BR INSENSITIVE2 (BIN2) promotes and restricts stomatal asymmetric cell division (ACD) depending on its subcellular localization, which is regulated by the stomatal lineage-specific scaffolding protein POLAR. BRs inactivate BIN2, but how they govern stomatal development remains unclear. Mapping the single-cell transcriptome of stomatal lineages with either exogenous BRs or the specific BIN2 inhibitor revealed that the two modes of BR activation triggered spatiotemporally distinct transcriptional responses. We established that when in a complex with POLAR and its closest homolog POLAR-LIKE1, BIN2 is insulated from BR-mediated inactivation, nevertheless, it remains sensitive to the inhibitor. Subsequently, BR signalling is attenuated in ACD precursors, whereas it remains active in epidermal cells that would undergo differentiation. Our study demonstrates how scaffold proteins contribute to cellular signal specificity of hormonal responses in plants.

---

The steroidal phytohormones, brassinosteroids (BRs), are essential regulators of growth and development and their signalling is one of the best-defined signal transduction pathways in plants^1^. The *Arabidopsis thaliana* SHAGGY/GSK3-like kinase 21 (*At*SK21)/BR-INSENSITIVE2 (BIN2) is a key negative regulator of BR signalling, because it inactivates two master transcription factors, BRASSINAZOLE RESISTANT1 (BZR1) and BRI1-EMS-SUPPRESSOR1 (BES1)/BZR2 via phosphorylation^2, 3^. BR binding to the plasma membrane (PM)-localized BR receptor complex initiates a signalling cascade that leads to BIN2 inactivation and degradation through dephosphorylation by the phosphatase BRI1 SUPPRESSOR1 (BSU1) and ubiquitination by the F-box protein KINK SUPPRESSED IN BZR1-1D (KIB1), respectively^4, 5^. Consequently, dephosphorylated BZR1 and BES1 translocate to the nucleus and regulate expression of target genes^6, 7^.

BIN2 is also an important regulator of the stomatal signalling pathway^8–11^. In Arabidopsis, the stomatal lineage is initiated from an undifferentiated meristemoid mother cell (MMC) that undergoes an asymmetric cell division (ACD), producing a small meristemoid and a large stomatal lineage ground cell (SLGC). The meristemoid either differentiates into a guard mother cell (GMC) that divides symmetrically and forms stomata or undergoes several amplifying ACDs to produce additional SLGCs^12^. The SLGCs give rise to pavement cells and new satellite meristemoids as a result of ACDs that are strictly controlled by the activity of the transcription factor SPEECHLESS (SPCH). SPCH is negatively regulated by a canonical mitogen-activated protein kinase (MAPK) signalling module that functions downstream of a ternary receptor-peptide complex, including members of the ERECTA family (ERf), the receptor-like protein TOO MANY MOUTHS (TMM) and their ligands, the secreted cysteine-rich peptides, EPIDERMAL PATTERNING FACTORs (EPFs)^13^. An intrinsic polarity complex, including the scaffolding proteins BREAKING OF ASYMMETRY IN THE STOMATAL LINEAGE (BASL), BREVIS RADIX-LIKE2 (BRXL2), and POLAR LOCALIZATION DURING ASYMMETRIC DIVISION AND REDISTRIBUTION (POLAR), is necessary for the ACDs^14–17^. BIN2 inhibits SPCH and the PM-localized MAPK components through phosphorylation and, consequently, limits or promotes stomatal development^8–10^. These two BIN2 activities are regulated by the scaffolding function of POLAR that anchors BIN2 to the PM, attenuating the MAPK signalling and enabling SPCH to drive ACDs in the nucleus^11^. Despite recent knowledge on the function of BIN2 in stomatal development^8–11^, the role of BRs in this process remains poorly understood^8, 9^.

By generating and exploiting a high-resolution single-cell gene expression map of the stomatal lineage in Arabidopsis, we found that BR signalling activation by either exogenous BRs or the plant-specific BIN2 inhibitor, bikinin^18^, triggered differential transcriptional responses with contrasting stomatal development outputs. Bikinin strongly suppressed stomata formation, whereas exogenous BRs induced divisions in the stomatal lineage. We established that this opposite effect was due to different BIN2 inactivation modes, while exogenous BRs acted via the BRI1-dependent signal transduction pathway to inactivate BIN2, bikinin directly inhibited it. When in a complex with POLAR and POLAR-LIKE1 (PL1) in the PM, BIN2 activity was enhanced and insulated from the BR-mediated inactivation, but it remained highly sensitive to bikinin. Monitoring the dynamics of the BES1 nuclear accumulation in stomatal lineage revealed that the BR signalling was downregulated in POLAR- and PL1-expressing ACD precursors, but that it was restored in SLGCs.

In summary, by understanding the molecular basis that determines the contrasting effects of exogenous BRs and bikinin on stomatal lineages, we provide a mechanistic insight into an intrinsic developmental program regulation in which cell-specific responses are generated through scaffolding molecules. We show that stomatal ACDs require BR signalling attenuation that is achieved via insulation of the BIN2 activity by POLAR and PL1.

## Results

### BRs and bikinin inversely affect stomatal development in Arabidopsis

Pharmacological inhibition of BIN2 with bikinin^18^ reduced stomatal density in the abaxial cotyledon and leaf epidermis of wild type Arabidopsis and suppressed stomatal clustering in the *bin2-1* and *bsu-q* mutants^9^. In contrast, the fact that wild type plants grown on exogenous BRs did not exhibit any epidermal phenotypes in cotyledons^8, 19^ was difficult to reconcile with the BR signalling activation by both bikinin and BRs^4, 18^. As BRs do not undergo long-distance transport^20^ and treatments on agar media^8, 19^ possibly limit the access of BRs to the above-ground tissues, wild type Arabidopsis seeds were germinated and grown for 3 days in liquid medium supplemented with increasing concentrations of either brassinolide (BL), the most active BR, or bikinin (Extended Data Fig. 1a,b). Exogenous BL significantly increased the total epidermal cell density in a concentration-dependent manner, because of an increased number of small (<200 μm^2^) non-stomatal cells, indicating an induction of additional cell divisions (Extended Data Fig. 1a,b). In particular, the small (<200 μm^2^) non-stomatal epidermal cells retained a low degree of asymmetry after the division (Extended Data Fig. 1c), calculated according to the cell size of the daughter cell after division^11, 19^. In contrast, and as previously reported^9^, bikinin strongly reduced the total epidermal cell density, mainly due to a reduction in the small (<200 μm^2^) non-stomatal cell density and in the stomatal index (Extended Data Fig. 1a,b). Altogether, these data indicate that BL and bikinin have an opposite impact on the BIN2 activity in stomatal lineage, although both activate the BR signaling^18^.

### BRs and bikinin trigger spatiotemporally distinct transcriptional responses

To capture the differences between exogenous BL and bikinin at the cellular level, we applied single-cell transcriptomics of the stomatal lineage. First, a high-resolution single-cell RNA-sequencing (scRNA-seq) map of stomatal lineage cells was generated as a control dataset (Fig. 1). For the profiling of stomatal lineage cells only, a fluorescence-activated cell sorting (FACS) was utilized on protoplast cells isolated from the aerial parts (cotyledons) of Arabidopsis seedlings that harboured the TMM reporters, *TMMpro:TMM-GFP* or *TMMpro:GFP*^21^ 5 days post germination (dpg) (Fig. 1a). The *TMMpro:GFP* protoplasts were derived from a DMSO-treated tissue (a mock control) for 2 h in liquid medium. After quality control and stringent filtering steps (see Methods for details), we retained 8005 and 4937 individual high-quality cells from these two independent experiments with a minimum UMI count of 2001 and 1481 (Fig. 1b), respectively. Plotting the transcriptomes from the two datasets by means of the Uniform Manifold Approximation and Projection (UMAP)^22^ revealed largely overlapping cell distributions between the translational and transcriptional TMM reporters used to generate these datasets, with the latter displaying a broader distribution range, consistent with its larger expression domain (Fig. 1a, b). Ten clusters reflecting the developmental transitions in the stomatal lineage were annotated with known stomatal markers^23–27^ (Fig. 1c-e and Supplementary Data 1). Overall, the UMAP topology was similar to the previously published single-cell resolution map of the stomatal lineage^23^, with the exception of an additional cell cluster with unknown cell identity. To validate the predictions from our control dataset, we generated 21 reporter lines of previously unknown differentially expressed genes (DEGs) selected from each cell cluster. Except for two genes, *AT3G48520* and *AT4G04840* chosen as DEGs in the SLGC and pavement cell clusters, respectively, all examined reporter lines perfectly recapitulated the predicted expression domains (Extended Data Fig. 2), confirming the cluster annotation.

**Fig. 1:**
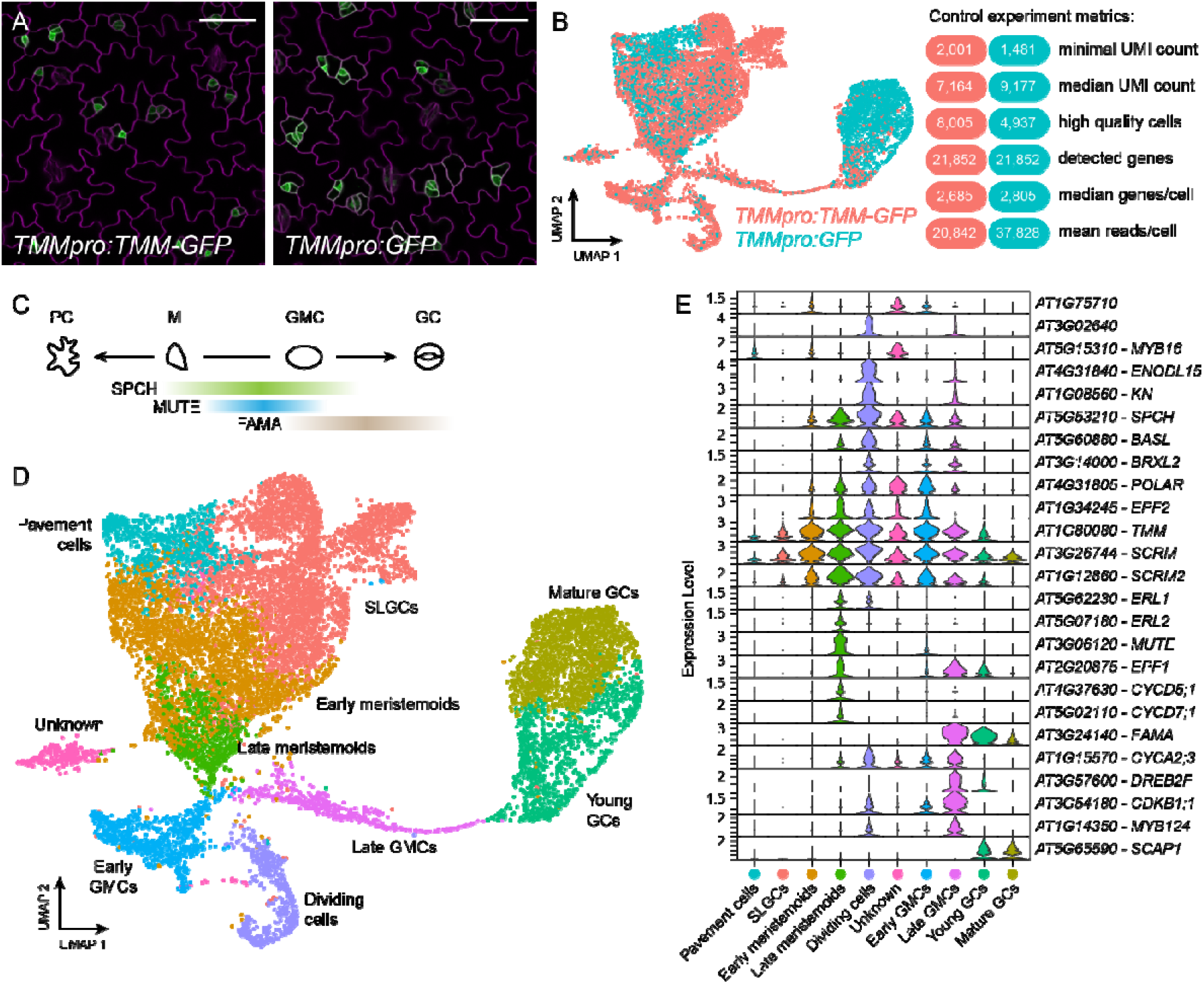
Single-cell RNA-sequencing of Arabidopsis stomatal lineage cells. **a,** Confocal images of the abaxial cotyledon epidermis of plants expressing the translational (*TMMpro:TMM-GFP*) and transcriptional (*TMMpro:GFP*) TMM reporters at 5 days post germination used for single-cell RNA-sequencing (scRNA-seq). Cell outlines (magenta) were marked with propidium iodide. Scale bar, 50 μm. **b,** UMAP visualization of integrated scRNA-seq datasets derived from *TMMpro:TMM-GFP* and *TMMpro:GFP*. Each dot represents an individual cell. Cell transcriptomes generated from each individual experiment are indicated by different colours. **c,** Schematic representation of stomatal development and expression levels shown by colour intensity of three key stomatal regulators, the transcription factors SPEECHLESS (SPCH), MUTE, and FAMA. **d,** Color-coded UMAP plot showing the classification of single cells into ten distinct clusters corresponding to different cell identities in the stomatal lineage. SLGC, stomatal lineage ground cell; GMC, guard mother cell; GC, guard cell. **e,** Violin plots depicting level (height) and proportion of cell expression (width) of known stomatal lineage marker genes across each cell cluster. Colours correspond to cell clusters as shown in **d**.

Next, we profiled sorted protoplasts derived from the aerial parts (cotyledons) of Arabidopsis seedlings expressing *TMMpro:GFP* at 5 dpg treated with BL (1 μM) and bikinin (50 μM) for 2 h in liquid medium. After filtering for high-quality cells, 7,011 and 5,909 cells were retained from the BL- and bikinin-treated samples, respectively, with a minimum UMI count of 591 and 606 (Fig. 2a). The two datasets were then merged with the DMSO-treated *TMMpro:GFP* control sample, which was generated at the same time and the Seurat package was used to transfer cell annotation labels to the combined dataset (Fig. 2b). The three samples were interspersed across the UMAP plot (Fig. 2b) and contributed equally to each cell cluster (Extended Data Fig. 3a), suggesting that the new dataset provides a good predictive power and that it can be used for further analysis of the spatiotemporal gene expression during stomatal development.

**Fig. 2:**
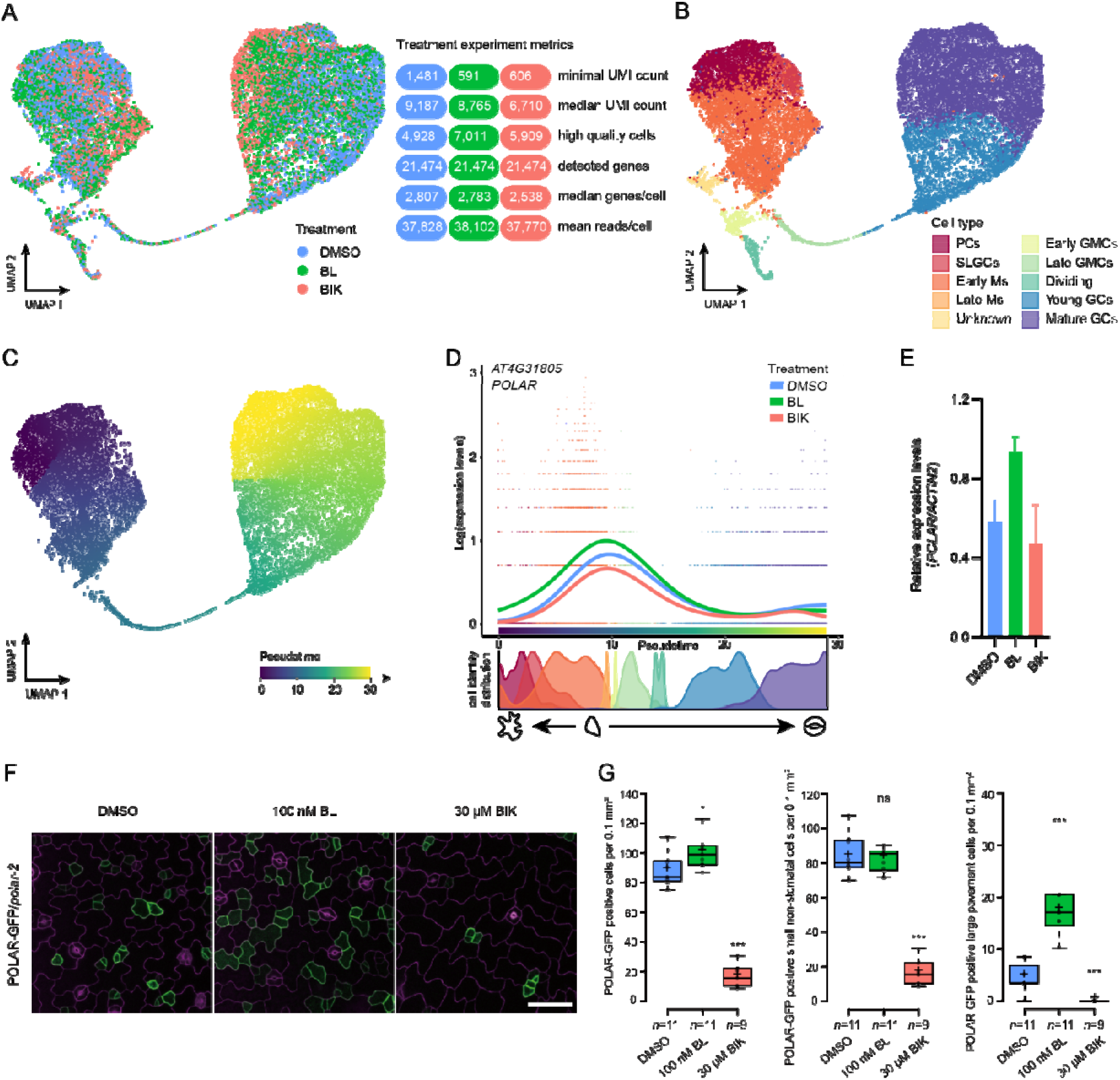
BRs and bikinin trigger spatiotemporally distinct transcriptional responses. **a,** UMAP visualization and experiment metrics of the integrated single-cell RNA-sequencing datasets derived from the aerial parts (cotyledons) of seedlings expressing *TMMpro:GFP* treated with brassinolide (BL, 1 μM), bikinin (BIK, 50 μM), and DMSO (mock) for 2 h in liquid medium. Each dot represents an individual cell. Cell transcriptomes are color-coded according to treatments. **b,** UMAP plot of the merged data with indications of the cell type assignment. **c,** Pseudotime assignment in colour code for the merged data. **d,** Expression trends of POLAR along pseudotime of the stomatal lineage trajectory after treatment with BL, bikinin, and DMSO (top) and distribution of the cell identities over the pseudotime trajectory, with colours denoting cell types as shown in **b** (bottom). **e,** Quantitative real-time PCR analysis of *POLAR*. Total RNA was extracted from 3-day-old wild type Arabidopsis seedlings treated as in **a**. Transcript levels were normalized to *ACTIN2* (AT3G18780). Five independent experiments with three technical replicates each were analysed. **f,** Expression of POLAR-GFP [*POLARpro:gPOLAR-GFP/polar-2* (line #14.5)] in response to BL and bikinin. Seeds were germinated and grown in liquid medium containing 100 nM BL, 30 μM bikinin, and DMSO (mock) for 3 days. Cell outlines (magenta) were marked with propidium iodide. Scale bar, 50 μm. **g,** Quantification of **f.** POLAR-GFP-positive cells calculated as small (<200 μm^2^) non-stomatal cells and large pavement cells (≧200 μm^2^) per 0.1 mm^2^. Data points are plotted as dots. Center lines show the medians. Crosses represent sample means. Box limits indicate the 25th and 75th percentiles as determined by R software. Whiskers extend 1.5 times the interquartile range from the 25th and 75th percentiles. Significant differences were determined with a single factor ANOVA. \*\*\**P*< 0.001, \**P*< 0.05. ns, no significant difference. *n*, number of cotyledons analysed.

To compare expression levels between treatments and across developmental transitions, we used the slingshot algorithm for trajectory inference^28^, thus determining a pseudotime value for every cell (Fig. 2c). When the trajectory was mapped to the cell clusters, a dual trajectory emerged, one from meristemoid-to-pavement (pseudotime 8 to 0) cells and one from meristemoid cells (pseudotime 8) through GMCs (pseudotime 12) into GCs (pseudotime 30). Thus, our dataset captured the entire developmental program of the stomatal lineage in pseudotime (Fig. 2c). Next, expression levels were calculated for each gene over the entire trajectory by means of the tradeSeq algorithm^29^, resulting in expression profiles that can be compared between the different treatments for each developmental moment (see Methods for details) (Supplementary Data 2). Then, we aimed at validating a number of genes that were predicted to be significantly differentially expressed along the pseudotime after BL or bikinin treatment with a focus on genes with known functions in stomatal development (Extended Data Fig. 3b-d and Supplementary Data 2). BL mostly upregulated stomatal markers expressed alongside the trajectory meristemoids-late meristemoids-early GMCs-GMCs, including *BASL, POLAR, EPF1, MUTE*, and the newly validated DEGs *AT5G11550* and *AT1G51405* for cluster “Early Ms” and “Late GMCs”, respectively (Fig. 2d and Extended Data Fig. 2a,c, 3b). In contrast, bikinin either had no effect (*BASL, EPF1, MUTE, AT5G11550*, and *AT1G51405*) or downregulated genes (*POLAR, EPF2, ERL1, CYCD7;1*,and the newly validated DEG for cluster “Early Ms”*AT3G62070*) in this trajectory (Fig. 2d and Extended Data Fig. 2a, 3b,c). Interestingly, bikinin upregulated the GMC/GC marker genes, *SIAMESE-RELATED4* (*SMR4*)^30^, *CYCLIN A2;3* (*CYCA2;3*)^27^, and genes expressed in GCs, such as *AT3G47675* (Extended Data Fig. 2d), *HIGH LEAF TEMPERATURE1* (*HT1*^31^, *K+ CHANNEL ARABIDOPSIS THALIANA2* (*KAT2*)^32^ (Extended Data Fig. 3d). Quantitative real-time PCR analysis (qRT-PCR) of either wild type seedlings (Fig. 2e) or *TMMpro:GFP*-positive protoplasts isolated by FACS from seedlings treated with BL, bikinin, and DMSO as done for the scRNA-seq work (Extended Data Fig. 3e), together with imaging and quantification of transcriptional reporters after similar treatments (Extended Data Fig. 3f,g) validated the expression patterns of *POLAR, BASL, EPF2, MUTE, SMR4*, and *CYCA2;3*. In addition, long-term treatments also confirmed that exogenous BRs promoted the expression of stomatal regulators, whereas bikinin downregulated them (Fig. 2f,g and Extended Data Fig. 4a-h). Taken together, these results reveal that exogenous BL and bikinin trigger spatiotemporally distinct transcriptional responses in stomatal lineage cells, possibly the reason for the observed different epidermal phenotypes.

### Constitutive overexpression of POLAR and PL1 leads to BR insensitivity

When examining the list of genes with significant differential responses to bikinin and BL (Supplementary Data 2), we noticed the known regulator of stomatal development and a direct interactor of BIN2, POLAR^11, 16^. POLAR was upregulated by BL and downregulated by bikinin from the early meristemoid to the late GMC stages (Fig. 2d-g). Consistently with the observed phenotypes (Extended Data Fig. 1) and the scRNA-seq data, bikinin suppressed the expression of POLAR-GFP in *POLARpro:gPOLAR-GFP/polar*^11^, whereas exogenous BL extended the POLAR-GFP expression mostly in the SLGCs. (Fig. 2f, g). As *POLAR* functions redundantly with its closest homolog, *PL1* (At5g10890)^11, 24^, we also examined the behavior of PL1-GFP (*PL1pro:gPL1-GFP*/Col-0) in the stomatal lineage (Extended Data Fig. 5). In the abaxial cotyledon epidermis of Arabidopsis, PL1-GFP polarized before the ACD in a manner similar to that of POLAR (Extended Data Fig 5a,b and Supplementary Video 1). PL1-GFP co-purified with BASL-mCherry, POLAR-mCherry, and BIN2-mCherry in tobacco (*Nicotiana benthamiana*) leaf epidermis (Extended Data Fig. 5c) and BASL-BFP efficiently polarized PL1-GFP, whereas the PL1-GFP expression alone lacked polarity (Extended Data Fig. 5d,e). As previously reported for POLAR^11^, in the presence of BFP-BASL, PL1-GFP excluded BIN2-mCherry from the nucleus and subsequently tethered it to the PM in a polar fashion (Extended Data Fig. 5f). Moreover, PL1 and POLAR behaved similarly under prolonged BL and bikinin treatments (Fig. 2f,g and Extended Data Fig. 4d). Altogether, these findings indicate that POLAR and PL1 probably function as part of the same polarity complex and that they might, at least in part, be accountable for the differential responses to BRs and bikinin observed in the stomatal lineage. In agreement, when treated with BL, the sensitivity of the double *polar pl1* mutant^11^ was lower than that of the wild type in terms of small (<200 μm^2^) non-stomatal cell density increase, whereas the impact of bikinin was equal to that of the wild type (Extended Data Fig. 1a,d).

To further test our hypothesis, we generated transgenic lines expressing the genomic (g) fragments of POLAR and PL1 driven by the *CaMV 35S* promoter [*35Spro:gPOLAR* (POLAR-OE) and *35Spro:gPL1* (PL1-OE)] (Fig. 3a,b and Extended Data Fig. 6a). Constitutive overexpression of either POLAR or PL1 in Arabidopsis resulted in an excess of small (<200 μm^2^) non-stomatal cells in the abaxial cotyledon epidermis at 3 dpg (Extended Data Fig. 6b,c), but the phenotype was milder than that of POLAR, when overexpressed in its native domain^11^. Moreover, these plants had more compact rosettes with dark-green and rounded leaves (Fig. 3a,b), resembling a *bin2-1* mutant or plants overexpressing BIN2^33^, which are defective in BR signalling. Root growth inhibition assays with increasing BL concentrations revealed that all overexpression lines were insensitive to exogenous BRs (Fig. 3c,d). In line with these observations, overexpression of *POLAR* or *PL1* also increased the phosphorylated and total BES1 protein levels (Fig. 3e,f). Consequently, BL treatment (100 nM) of POLAR-overexpressing plants did not efficiently induce BES1 dephosphorylation as in the wild type, even more prominently so with PL1 overexpression (Fig. 3e,f). In contrast, treatment with bikinin (50 μM) completely dephosphorylated BES1 in both overexpression lines, similarly to the wild type (Fig. 3e,f). Thus, when in a complex with POLAR or PL1, the kinase activity of BIN2 was efficiently inhibited by bikinin, but not by BL. Nevertheless, BIN2 in a complex with either POLAR or PL1, was still able to phosphorylate BES1 without leading to its turnover. Overall, we conclude that BIN2, and possibly its homologs, do not undergo BR-mediated inactivation, but remain sensitive to bikinin when in a complex with POLAR and PL1.

**Fig. 3:**
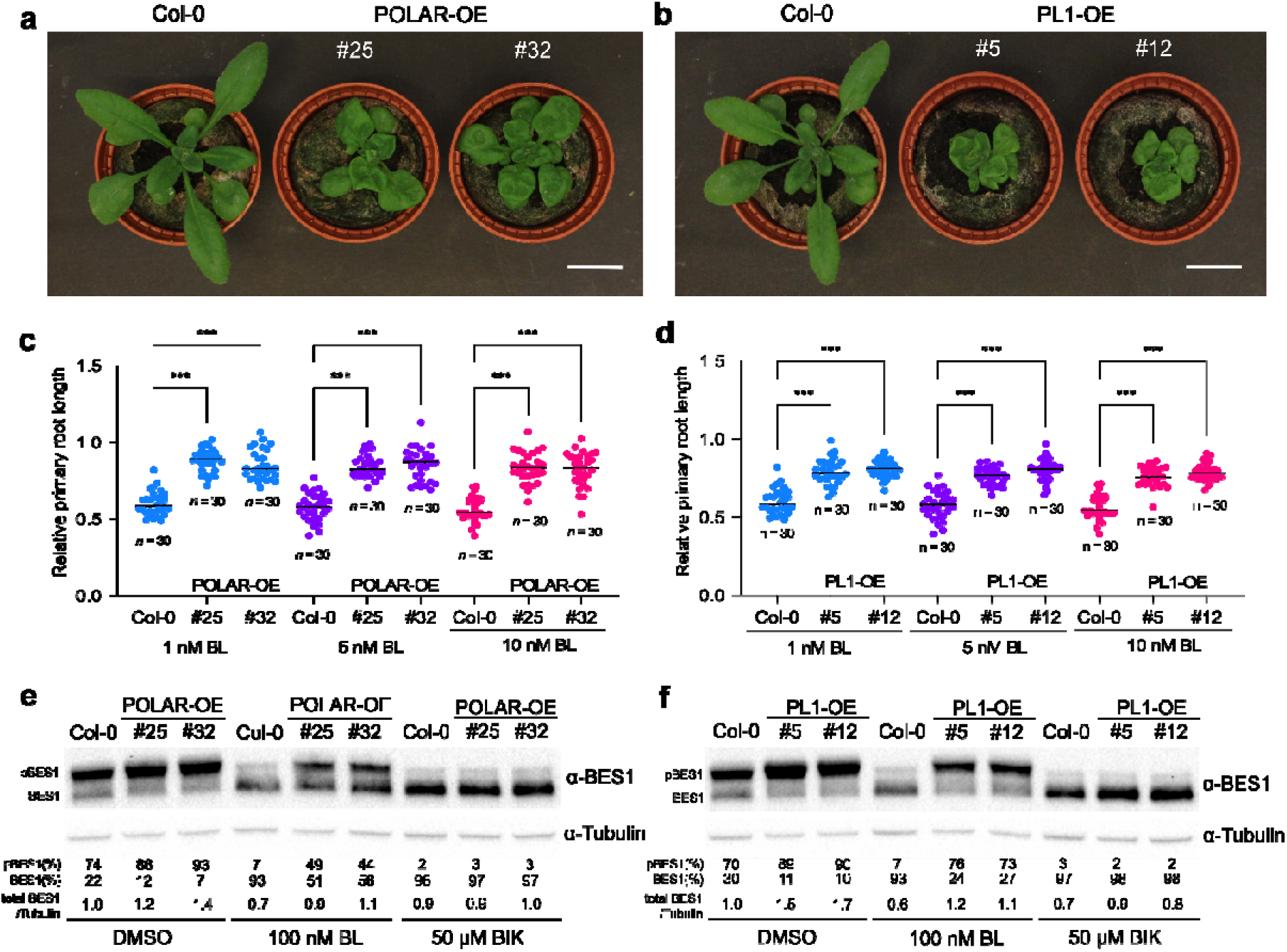
Overexpression of POLAR and POLAR-LIKE1 (PL1) led to BR insensitivity. **a,b,** Phenotypes of 4-week-old wild type (Col-0) Arabidopsis and two independent homozygous transgenic lines overexpressing POLAR [*35Spro:gPOLAR* (POLAR-OE), lines #25 and #32] (**a**) and PL1 (*35Spro:gPL1*, PL1-OE, lines #5 and #12) (**b**). Scale bars, 2 cm. **c,d,** Root growth inhibition assay for testing BR sensitivity of Col-0, POLAR-OE (lines #25 and #32) (**c**), and PL1-OE (lines #5 and #12) (**d**) plants. Plants were grown in the presence of increasing BL concentrations (1 nM, 5 nM, and 10 nM) and DMSO (mock) for 5 days. The primary root length in the presence of BL is relative to the DMSO control. All individual data points are shown and black horizontal bars represent the means. *n*□, □ number of roots analysed. Asterisks highlight significant differences from Col-0 by two-way ANOVA with Tukey’s multiple comparisons test (\*\*\**P*< 0.001). The data are representative of three independent experiments. **e,f,** Endogenous BES1 levels in Col-0, POLAR-OE (lines #25 and #32) (**e**), and PL1-OE (lines #5 and #12) (**f**) plants after treatment with BL and bikinin (BIK). Five-day-old Arabidopsis seedlings were transferred to liquid media supplemented with 100 nM BL for 2 h or 50M bikinin for 1 h. Quantification shows the ratio between phosphorylated (pBES1) and unphosphorylated BES1 (%). The total BES1 abundance is relative to the DMSO-treated control. The data shown are representative of three independent experiments.

### POLAR and PL1-dependent scaffolding of BIN2 increases its stability and activity

To investigate the cause for the BR insensitivity of POLAR and PL1-overexpressing plants, we introduced each *35Spro:gPOLAR* or *35Spro:gPL1* construct into the *BIN2pro:gBIN2-GFP/Col-0* (line #32.7)^11^ ensuring that the BIN2-GFP levels remain the same (Fig. 4a and Extended Data Fig. 7a,b). In parallel, we also transformed the *POLARpro:gPOLAR-mCherry* and *PL1pro:gPL1-mCherry* constructs into the same BIN2-GFP line *[BIN2pro:gBIN2-GFP*/Col-0 (line #32.7)] (Extended Data Fig. 7c-e). Examination of the abaxial cotyledon epidermis by confocal microscopy revealed a strong accumulation of the BIN2-GFP signal when either POLAR or PL1 were overexpressed (Fig. 4a and Extended Data Fig. 7c,d). As previously reported for POLAR^11^, overexpression of PL1-mCherry in its endogenous domain resulted in a strong overproduction of small stomatal lineage cells (Extended Data Fig. 7d). BIN2-GFP accumulated as a result of an overall increase in protein abundance, because no changes in BIN2 transcripts were detected (Extended Data Fig. 7b,e), as also confirmed by western blot analysis of BIN2-GFP in POLAR-OE and PL1-OE plants with α-GFP antibodies (Fig. 4b). Notably, the BIN2 accumulation was higher in PL1-OE plants, consistent with the higher PL1 amounts, than that with POLAR in the POLAR-OE plants (Fig. 4b and Extended Data Fig. 7b), hinting at a dose-dependent effect. In addition to its stabilization, the activity of BIN2 also increased, as shown with an antibody recognizing the phosphorylated threonine in the activation loop of the animal GSK3 (α-pTyr279-216-GSK3α/β) and frequently used to detect an active BIN2 in plants^5^ (Fig. 4b). Application of BL only slightly reduced the BIN2 protein and kinase activity levels and did not completely dephosphorylate BES1 (Fig. 4b), implying an increased BIN2 abundance and as a result activity, when POLAR or PL1 are overexpressed. Consistent with these results, nuclear fractionation experiments with BIN2-GFP-expressing lines in the wild type, POLAR-OE, and PL1-OE backgrounds revealed that endogenously phosphorylated BES1 was more abundant in the cytoplasm of POLAR-OE or PL1-OE plants than that of the control (Fig. 4c). Exogenous BL (100 nM) fully dephosphorylated BES1 in the nucleus of BIN2-GFP plants, whereas the nuclear BES1 pool in POLAR-OE or PL1-OE plants was sensitive to BL, albeit to a lesser extent. The cytoplasmic pool of BES1 was reduced in the presence of BL in the control, whereas in the POLAR-OE or PL1-OE plants, BES1 remained insensitive to the hormone (Fig. 4c). Altogether, these observations prompted us to examine whether BES1 is recruited to the POLAR and PL1-BIN2 complex. Indeed, co-immunoprecipitation assays in Arabidopsis (Fig. 4d) revealed that BES1 co-purified with POLAR, PL1, and BIN2.

**Fig. 4:**
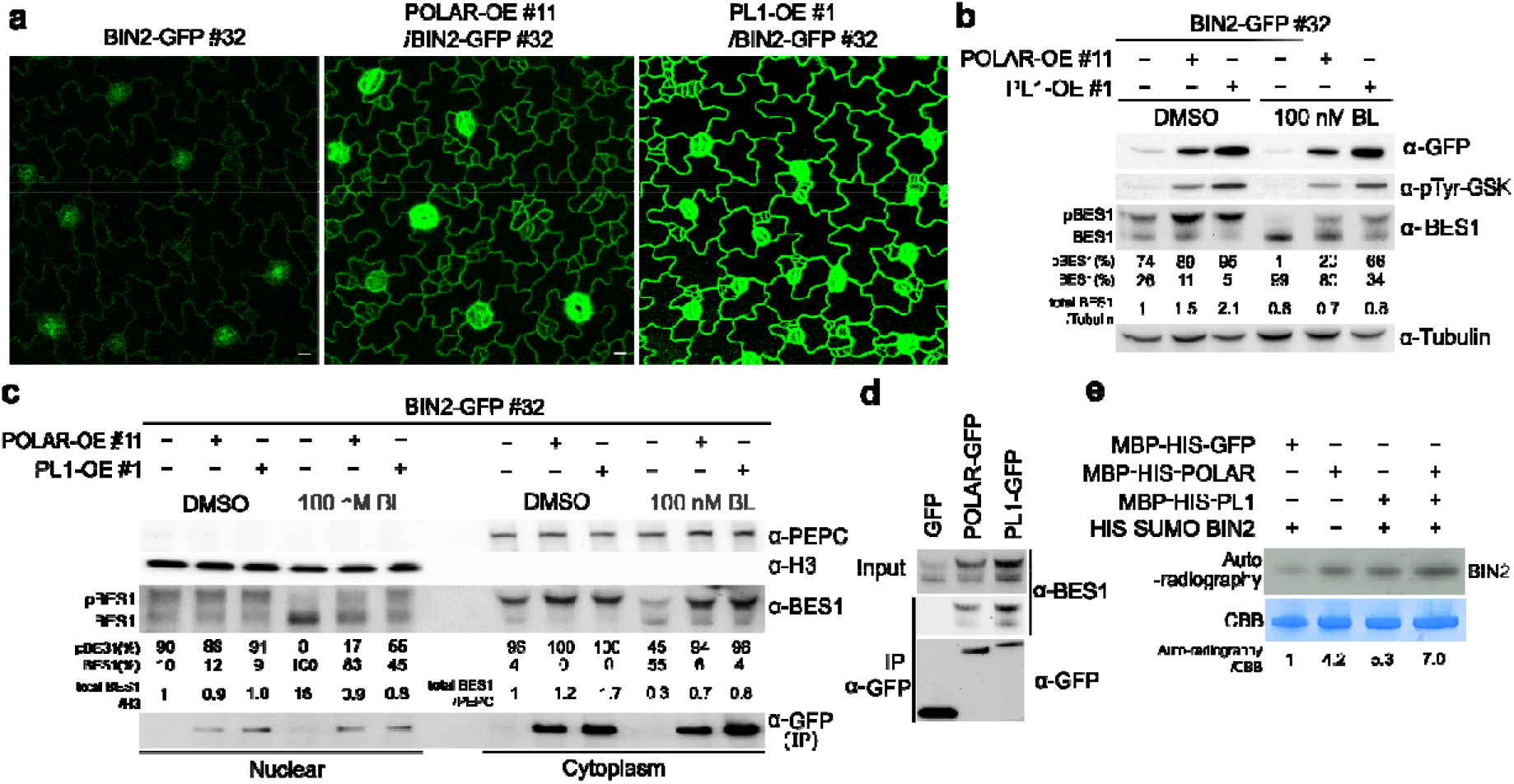
Scaffolding of BIN2 by POLAR and PL1 increased its stability and activity. **a,** Expression pattern of BIN2-GFP in wild type [*BIN2pro:gBIN2-GFP*/Col-0 (line #32.7)], POLAR-OE *[35Spro:gPOLAR/BIN2pro:gBIN2-GFP* (line #32.7)/Col-0 (line #11)], and PL1-OE *[35Spro:gPL1/BIN2pro:gBIN2-GFP* (line #32.7)/Col-0 (line #1)] backgrounds. The abaxial cotyledon epidermis of Arabidopsis seedlings was observed by confocal microscopy at 3 days post germination (dpg). Scale bars, 20 μm. **b,** Increase in protein levels of BIN2-GFP when POLAR or PL1 were overexpressed. Homozygous Arabidopsis seedlings expressing BIN2-GFP as in **a** were treated with 100 nM brassinolide (BL) and DMSO (mock) for 1 h in liquid medium. BIN2-GFP was captured by immunoprecipitation and detected with α-GFP and α-pTyr-GSK antibodies. Tubulin was used as a loading control and detected with α-tubulin antibody. Endogenous BES1 levels were detected with α-BES1 antibodies. Quantification of the ratio unphosphorylated/phosphorylated (p)BES1 was indicated in %. The total BES1 abundance is relative to the DMSO-treated BIN2-GFP control. **c,** Nuclear fractionation of Arabidopsis seedlings expressing BIN2-GFP as in **a** and treated with 100 nM BL as in **b**. Phosphoenolpyruvate carboxylase (PEPC) and histone H3 (H3) were used as a cytoplasmic marker and a nuclear marker, respectively. BIN2-GFP was detected after immunoprecipitation with α-GFP antibody. Endogenous BES1 levels were detected with α-BES1 antibody. IP, immunoprecipitation. The blot is labelled as in **b**. The total BES1 protein abundance was quantified relative to H3 and to PEPC. **d,** BES1 co-immunoprecitated with POLAR-GFP and PL1-GFP in Arabidopsis seedlings at 3 dpg. Transgenic POLAR-GFP-OE [*35Spro:gPOLAR-GFP*/Col-0 (line #3.1)], PL1-GFP-OE [*35Spro:gPL1-GFP*/Col-0 (line #1.6)], and GFP-OE [*35Spro:GFP*/Col-0] plants were used. GFP-OE was used as a negative control. The α-GFP antibody was used for immunoprecipitation and the blots were detected with α-GFP and α-BES1. **e,** *In vitro* autophosphorylation assay of BIN2 in the presence of POLAR and PL1. SUMO-HIS-BIN2 was incubated with MBP bead-bound MBP-HIS-GFP, MBP-HIS-POLAR, and MBP-HIS-PL1. After the reaction with 32P-γ-ATP for 2 h at 30 °C, only the supernatant fraction containing SUMO-HIS-BIN2 of each sample was collected and separated by SDS-PAGE. Autoradiography signal shows kinase activity. Quantification shows the autophosphorylation activity of SUMO-HIS-BIN2 relative to its protein level. CBB, Coomassie brilliant blue.

To explore whether POLAR or PL1 affected the BIN2 kinase activity through their direct interaction, we examined BIN2 autophosphorylation in vitro in the presence of POLAR or PL1 (Fig. 4e). BIN2 kinase activity increased in the presence of POLAR or PL1, but more importantly when the two scaffolds were added simultaneously. Collectively, we show that when in a complex with POLAR and PL1 in the PM, BIN2 is stabilized and its kinase activity is increased, consequently leading to BR insensitivity.

### BR signalling is attenuated in ACD precursors

Given that POLAR and PL1 are exclusively expressed in the early stomatal lineage^11, 16, 24^ (Extended Data Fig. 5a,b and Supplementary Video 1), we assumed that BIN2 insulation from BR-mediated inactivation in these cells is important for stomatal development. To gain more insight into the BR signaling regulation in stomatal lineage, we analyzed the BES1-GFP^34^ localization in the abaxial cotyledon epidermis of wild type Arabidopsis plants expressing BES1pro:gBES1-GFP^34^ and the PM marker *ML1pro:mCherry-RCI2A*^35^ at 2 dpg (Fig. 5a). The nuclear accumulation of BES1-GFP is routinely used as a readout for active BR signaling^3^. BES1-GFP was ubiquitously expressed throughout the cotyledon epidermis, but displayed differences in its nuclear localization in the stomatal lineage. BES1-GFP was enriched in the nuclei of SLGCs or pavement cells, but was less abundant in meristemoids or MMCs (Fig. 5a), consistent with the observed enhanced PM association of the endogenous BIN2 and its homologs in these cells^11^. A time-lapse imaging was performed over a 9-h course to track BES1-GFP before and after the amplifying ACD (Fig. 5b and Supplementary Video 2). Before the ACD, the nuclear BES1-GFP signal in meristemoids decreased, but after division, it increased in the nucleus of SLGCs, while remaining low in the nucleus of the newly formed meristemoid, albeit reaching the same expression levels as before the ACD (Fig. 5b). Examination of the co-localization of BES1-GFP (*BES1pro:gBES1-GFP*/Col-0) with SPCH-mCherry (*SPCHpro:gSPCH-mCherry*/Col-0) and mCherry-BASL (*BASLpro:mCherry-gBASL*/Col-0) before and after ACDs (Fig. 5c) revealed that, as anticipated, the BES1-GFP signal was strongly reduced in meristemoids with highly expressed SPCH and polarized BASL at the cell cortex before division (Fig. 5c), hinting at a attenuation of the BR signalling. In contrast, the nuclear signal of BES1-GFP in *polar pl1* double mutant remained enriched in meristemoids before and after the amplifying ACD (Fig. 5e and Extended Data Fig. 8a,b). Taken together, our data support that the ACD in meristemoids requires BR signalling attenuation, as substantiated by the expression of BR biosynthetic enzymes, of which a feedback inhibition via the BR signalling is known^36^. As predicted, DWF4-GFP (in *DWF4pro:DWF4-GFP/dwf4*)^20^ and GFP-BR6OX2 (in *BR6OX2pro:GFP-BR6OX2/br6ox1 br6ox2*)^20^ were enriched in meristemoids (Fig. 5d), thus supporting the notion that the BR signalling is downregulated in ACD precursors assigned to the BIN2 activity insulation by POLAR and PL1.

**Fig. 5:**
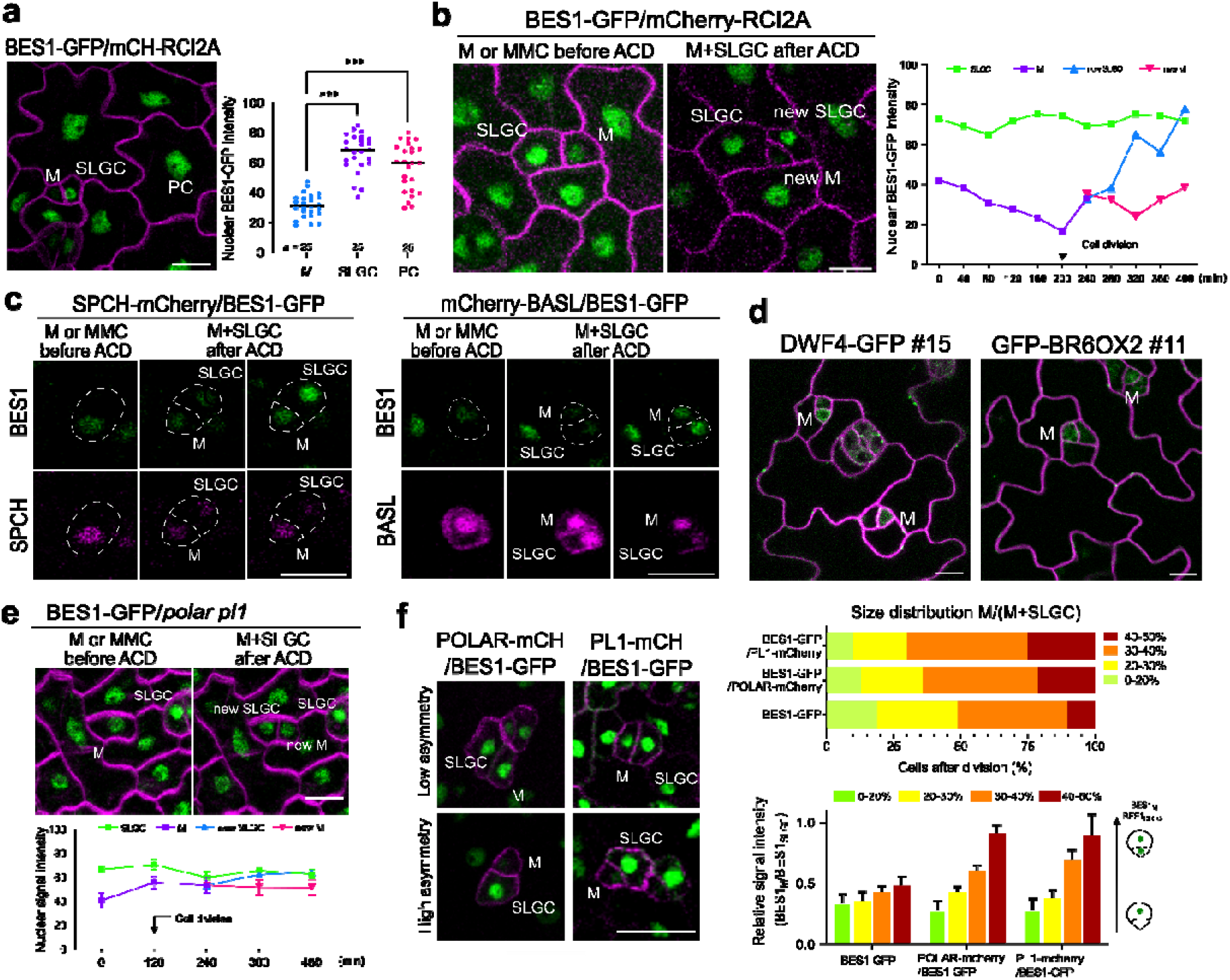
BR signalling is attenuated in ACD precursors. **a,** Localization of BES1-GFP (*BES1pro:gBES1-GFP/ML1pro:mCherry-RCI2A*/Col-0) in stomatal lineage in the abaxial cotyledon epidermis of Arabidopsis seedlings. Quantification shows the nuclear intensity of BES1-GFP in meristemoid (M), stomatal lineage ground cell (SLGC), and pavement cell (PC). All individual data points are shown and black horizontal bars represent the means. Asterisks highlight significant differences from meristemoids by one-way ANOVA and Dunnett’s multiple comparisons tests (\*\*\**P*< 0.001). *n*, number of cells analysed. **b,** Expression pattern of BES1-GFP before and after ACD. The nuclear BES1-GFP intensity was quantified over time as indicated. Arrow marks the time point of cell division. **c,** Expression pattern of BES1-GFP combined with either SPCH-mCherry *[SPCHpro:gSPCH-mCherry/BES1pro:gBES1-GFP* (line #5)] or mCherry-BASL [*BASLpro:mCherry-gBASL/BES1pro:gBES1-GFP* (line #9)] in stomatal lineage before and after ACD. Cell outlines were marked in dashed line. **d,** Expression pattern of DWF4-GFP [*DWF4pro:gDWF4-GFP/dwf4* (line #15.4)] and GFP-BR6OX2 [*BR6OX2pro:gBR6OX2-GFP/br6ox1 br6ox2* (line #11.5). **e,** Expression pattern of BES1-GFP in the *polar pl1* mutant [*BES1pro:gBES1-GFP/polar pl1* (line #11)] before and after ACD. The BES1-GFP nuclear signal intensity was quantified over time as indicated. Arrow marks the time point of cell division. **f,** Expression pattern of BES1-GFP in POLAR-OE [*POLARpro:gPOLAR-mCherry/BES1pro:gBES1-GFP*/Col-0 (line #3)] and PL1-OE [*PL1pro:gPL1-mCherry/BES1pro:gBES1-GFP*/Col-0 (line #2)]. The representative images show the expression of BES1-GFP in the cells with different degrees of asymmetry between meristemoid and SLGC after cell division. Top, quantification showing the size distribution between meristemoids and SLGCs after cell division in the wild type [*BES1pro:gBES1-GFP*/Col-0], POLAR-OE [*POLARpro:gPOLAR-mCherry/BES1pro:gBES1-GFP/Col-0* (line #3)], and PL1-OE [*PL1pro:gPL1-mCherry/BES1pro:gBES1-GFP/Col-0* (line #2)]. The area of each cells was measured and classified into four groups as indicated by the size distribution between meristemoid and SLGC after cell division (%). From each genotype, 100 meristemoids and 100 SLGCs were analysed. Bottom, quantification of the nuclear signal intensity of BES1-GFP in meristemoid relative to SLGC, presented according to the size distribution. Error bars represent SD. The abaxial cotyledon epidermis was examined by either time-lapse imaging 2 days post germination (dpg) in **b**, **c**, and **e** or by confocal microscopy at 3 dpg in **a**, **d**, and **f.** Scale bars, 10 μm. The experiments were repeated independently three times.

The scRNA-seq data and validation showed that exogenous BRs upregulate early stomatal regulators, consistently with the known stabilization of the SPCH protein by BRs^8^ due to the PM scaffolding and BIN2 insulation by POLAR and PL1 to allow MAPK inhibition, resulting in induction of ectopic epidermal cell divisions, a phenotype similar to that of overexpressed POLAR and PL1 in their native domains (Extended Data Fig. 7c,d). To assess the BES1-GFP nuclear localization in these overexpressing lines, we introduced the *POLARpro:gPOLAR-mCherry* and *PL1pro:gPL1-mCherry* constructs into the *BES1pro:gBES1-GFP*/Col-0 plants (Fig. 5f). The nuclear BES1-GFP intensity was classified according to the cell size after division^11, 19^. As previously reported for POLAR-GFP^11^, overexpression of PL1-GFP also reduced the asymmetry in the cell clusters, because most of the cells produced from meristemoid divisions had similar sizes (Fig. 5f). BES1-GFP had different fluorescence intensities depending on the level of cellular asymmetry after division when either POLAR or PL1 was overexpressed (Fig. 5f). BES1-GFP was strongly reduced in small cells with a high degree of asymmetry, whereas this pattern was not observed in cells with low asymmetry levels. As the stomatal index did not increase in the POLAR-overexpressing plants^11^, we conclude that cells with increased BES1 expression will probably acquire a pavement cell fate. In summary, exogenous BRs or overexpression of the scaffolds POLAR and PL1 in the stomatal lineage trigger ectopic cell divisions, alleviating SPCH inhibition by BIN2 in meristemoids and SLGCs as a consequence.

## Discussion

Understanding how different cell types interpret plant hormones to initiate cell type-specific responses remains a major challenge in plant developmental biology. Here, identification of the molecular mechanism behind the different effects of exogenous BR signalling activators (BL and bikinin) on stomatal lineages combined with single-cell transcriptomics and live imaging revealed the mechanistic framework underlying the involvement of BRs in the induction of stomatal ACDs. The role of BRs in stomatal development has long been a subject of debate, because both negative^9, 10^ and positive effects^8, 19, 37^ have been reported. Interestingly, a negative role relied mostly on the use of the small molecule bikinin^18^. Indeed, bikinin treatment suppressed divisions in stomatal lineage, whereas exogenous BRs promoted them. Although bikinin activates BR responses similarly as BRs and its application phenocopies plants with constitutive BR responses, some genes, among which stomatal regulators, have been reported to be controlled in an opposite manner^18^. These dissimilarities might lay in the different BIN2 inhibition mode. While bikinin acts directly at the BIN2 level^18^, BRs act through a signalling pathway mediated by protein-protein interactions^1^. Hence, we hypothesized that the BR-mediated BIN2 inactivation is regulated differently in stomatal lineages. This concept was further confirmed by sequencing the transcriptome of individual cells in stomatal lineages and by identifying the behaviour of each stomatal lineage cell type after treatment with either BL or bikinin. The pseudotime analysis revealed that most of the early stomatal lineage genes (*BASL, POLAR, EPF1*, and *MUTE*) along the meristemoid-to-GCs trajectory are upregulated by BRs, whereas they are either not affected (*BASL, EPF1*, and *MUTE*) or suppressed (*POLAR, EPF2, CYCD7;1*, and *ERL1*) by bikinin.

Interestingly, bikinin upregulated the cyclin-dependent kinase inhibitor SMR4 that was recently reported to repress ACDs through inhibition of the activity of CYCD7;1^30^, which is specifically expressed just prior to the symmetric GC-forming division^38^. Given that expression of key stomatal regulators, such as *POLAR, PL1, BASL*, and *MUTE* are induced by *SPCH*^15^, the prolonged expression of POLAR and PL1 in stomatal lineage caused by exogenous BRs implies that BR-induced ectopic cell divisions are triggered by SPCH stabilization^8^. Indeed SPCH protein levels were upregulated by BRs^8^ and downregulated by bikinin (Extended Data Fig. 4a). In view of the broadened role for SPCH in stomatal lineage^23^, BRs probably stabilize SPCH not only in meristemoids, but also in SLGCs and/or GCs. Therefore, the cell divisions induced by BRs might not all result in stomata. Exogenous BRs reduced the asymmetry of the cell divisions in a manner similar to that by overexpression of POLAR, which did not increase the stomatal index^11^. Thus, exogenous BRs positively regulated epidermal cell divisions, possibly through SPCH stabilization and consecutive upregulation of POLAR and PL1 that additionally reinforced the SPCH activity through BIN2 scaffolding.

Previously, POLAR had been shown to not only polarize, but also to enhance the PM-association of BIN2 and its homologs in ACD precursors^11^. The PM scaffolding of BIN2 by POLAR resulted in alleviation of the SPCH inhibition by BIN2 and MAPK, followed by ACD induction ^8–11^. A similar scaffolding mechanism for BIN2 was described in the protophloem of the Arabidopsis root, in which OCTOPUS (OPS) recruits BIN2 to the PM, thus preventing its inhibitory activity in the nucleus. Overexpression of OPS led to constitutive BR responses^39^, but overexpression of POLAR and PL1 caused BR insensitivity, while retaining sensitivity to bikinin. The BR resistance was caused by an increased BIN2 protein stability and kinase activity. Thus, when in a complex with POLAR and PL1, BIN2 is insulated from BR-mediated inactivation and degradation via a still unknown mechanism. Moreover, the PM-recruited and stabilized BIN2 remained active and accumulated phosphorylated BES1 in the cytoplasm, therefore restraining the BR signalling. The accumulation of dephosphorylated BES1 is used as a proxy for BR signalling activation^20, 34^. Analysis in the wild type revealed a strong reduction of the nuclear BES1 accumulation in ACD precursors, whereas the nuclear BES1-GFP was enriched in SLGCs and PCs, suggesting that BR signalling downregulation precedes the ACD. Taken together, we propose a model (Extended Data Fig. 8c), in which BIN2 is recruited to the PM by POLAR and PL1 in ACD precursors. Consequently, BIN2 is stabilized, hyperactive, and, hence, protected from BR-mediated inactivation. As a result, BR signalling in ACD precursors is attenuated to allow cell division via SPCH and to preclude cell differentiation. In contrast, in SLGCs, in the absence of POLAR and PL1 expression, BIN2 is inactivated by BRs to permit cell differentiation coinciding with an increase in nuclear BES1. Our work reveals that BR signalling regulated through BIN2 scaffolding and insulation ensures correct stomatal patterning in the leaf epidermis.

## Methods

### Plant materials and growth conditions

Seeds of *Arabidopsis thaliana* (L.) Heyhn. Columbia-0 (Col-0) and of the described genotypes sown on half-strength Murashige and Skoog (½MS) agar plates or in liquid ½MS medium without sucrose were stratified for 2 days in the dark at 4°C. The seeds were germinated and grown at 22 °C and under a 16-h light/8-h dark photoperiod for 3, 5, or 7 days, according to the experimental purposes. When indicated, brassinolide (BL) (OlChemIm, Ltd.) or bikinin (home synthesized) were added to the medium at a final concentration of 1, 5, 10, or 100 nM for BL, and 10, 30, or 50 μM for bikinin. The following transgenic Arabidopsis lines or mutants have been described previously: *TMMpro:TMM-GFP/Col gl1-1, TMMpro:GUS-GFP/Col gl1-1*^21^, *ML1pro:mCherry-RCI2A*/Col-0^35^, *POLARpro:gPOLAR-GFP/polar*, *BIN2pro:gBIN2-GFP*/Col-0, *POLARpro:gPOLAR-mCherry/BIN2pro:gBIN2-GFP*, and *polar pl1*^11^, *SPCHpro:SPCH-GFP/spch-3*^8^, *BES1pro:BES1-GFP*/Col-0^34^, *EPF1pro:erGFP*/Col-0^40^, *MUTEpro:MUTE-GFP/mute*^41^, *MUTEpro:GFP-GUS*^42^, *FAMApro:FAMA-GFP*/Col-0^43^, and *SMR4pro:GFP-GUS*/Col-0^44^. For the phenotypic analysis of 4-week-old Arabidopsis plants, 7-day-old seedlings grown in ½MS plates were transferred to soil and were grown at 21°C under a 16-h light/8-h dark regime. Wild type tobacco (*Nicotiana benthamiana*) plants were grown in the greenhouse at 25°C under a normal light regime (14 h light/10 h dark). For the single-cell sequencing experiment, Arabidopsis plants of *TMMpro:TMM-GFP/Col gl1-1* and *TMMpro:GUS-GFP/Col gl1-1*^21^ were grown vertically on solid ½MS plates without sucrose under continuous 24 h light conditions for 5 days.

### Generation of constructs and transgenic lines

The transcriptional or translational reporter constructs for validating the cell cluster annotation of the single-cell sequencing are listed in Supplementary Table 1. For *AT5G11550, AT2G41190, AT1G75900, AT1G04110, AT1G51405, AT4G04840*, and *AT3G08770*, the promoter region (up to ~3 kb) and the full-length genomic fragment of each gene were recombined with GFP into pB7m34GW by means of the Gateway LR clonase enzyme mix (Thermo Fisher Scientific). The other expression constructs were generated via Golden Gate assembly (NEB). The promoter region (up to ~3 kb) upstream of the transcriptional start was cloned into the pGGA000 entry vector, whereas the full-length genomic coding region was cloned into the pGGC000 entry vector^45^. For translational reporter constructs, entry clones bearing the promoter, the genomic coding region, or GFP were recombined into the final expression vector pFASTRK-AG^46^. For transcriptional reporter constructs, entry modules carrying the promoter region, a nuclear localization signal (NLS) linker, or GFP were cloned into pGGB-AG^45^. Primers are listed in Supplementary Table 2. The protocol for Golden Gate assembly has been described previously^46^. Of each entry plasmid 100 ng was combined with 1× Cutsmart buffer (NEB), 1 mM ATP, 10 units of BsaI (NEB), and 200 units of T4 DNA ligase (NEB). The Golden Gate assembly reaction was carried out under the following conditions: 20 cycles of 37°C for 2 min followed by 16°C for 2 min; 50 C for 5 min; and 80°C for 5 min. The constructs were transformed into Col-0 or *ML1pro:mCherry-RCI2A* plants.

The MultiSite Gateway system (Invitrogen) was used for the generation of the plasmid constructs. For *UBQ10pro:PIP2A-mCherry*, the coding sequence of *PLASMA MEMBRANE INTRINSIC PROTEIN 2A* (*PIP2A*) was cloned into pENTR/D-TOPO and was subsequently recombined with pDONR-p4p1r-UBQ10pro and pDONR-p2rp3-mCherry into pH7m34GW. pDONR-p2rp3-BASL^11^ was recombined with pDONR-p4p1r-BASLpro^11^ and pDONR221-GFP into pH7m34GW to generate *BASLpro:GFP-BASL*. The expression cassette of *UBQ10pro:PIP2A-mCherry* was then recombined with *BASLpro:GFP-BASL* by means of the Gibson assembly to generate the *BASLpro:GFP-BASL-UBQ10pro:PIP2A-mCherry* construct, which was transformed into Col-0 plants. To generate the *pEPF2pro:EPF2-GFP* construct, the *EPF2* genomic fragment containing 1737 bp of the promoter, three exons, and two introns was amplified by PCR and recombined into the pK7FWG construct. The construct was subsequently transformed into Col-0 plants. For *POLAR-LIKE1 (PL1)*, the genomic fragment containing 1766 bp of the promoter, two exons, and one intron was cloned into pDONR-p4p1r (primers are listed in Supplementary Table 2) and recombined with pDONR221-mCherry or pDONR221-GFP into the pB7m24GW,3 vector to generate the *PL1pro:gPL1-mCherry* or *PL1pro:gPL1-GFP* constructs, which were transformed into *BIN2pro:gBIN2-GFP*/Col-0^11^ and Col-0 plants to generate *PL1pro:gPL1-mCherry/BIN2pro:gBIN2-GFP*/Col-0 and *PL1pro:gPL1-GFP*/Col-0 transgenic plants, respectively. The *PL1pro:gPL1-mCherry* construct and the previously described constructs, including *POLARpro:gPOLAR-mCherry, SPCHpro:SPCH-mCherry*, and *BASLpro:mCherry-gBASL*^11^ were transformed into *BES1pro:gBES1-GFP*/Col-0 plants^34^. The genomic fragment of *BES1* containing 1563 bp of the promoter, 3 exons, and 2 introns was cloned into pDONR-p4p1r (primers are listed in Supplementary Table 2) and recombined with pDONR221-GFP into the pH7m24GW,3 vector to generate the *BES1pro:gBES1-GFP* construct. The construct was subsequently transformed into Col-0 and *polar pl* plants^11^. For the generation of overexpression lines, genomic DNA (gDNA) of *PL1* was cloned into pDONR221. gDNA of *PL1* and *POLAR*^11^ were recombined into the pK7WG2 vector containing the *Cauliflower mosaic virus (CaMV) 35S* promoter, of which the resulting constructs were transformed into Col-0 and *BIN2pro:gBIN2-GFP*/Col-0^11^, respectively. For transient expression in tobacco, the *35Spro:BIN2-GFP, 35Spro:TagBFP2-BASL, 35Spro:mCherry-BASL*, and *35Spro:POLAR-mCherry* constructs had been described previously^11^. cDNA of *PL1* was cloned into pDONR221 and recombined in pK7FWG2 containing the *35S* promoter and the C-terminal *GFP*-coding sequence to generate the construct *35Spro:PL1-GFP*. The *BIN2* cDNA^11^ was recombined with the *35S* promoter and *mCherry* into the pB7m34GW vector to generate *35Spro:BIN2-mCherry*.

For the purification of MBP-HIS-POLAR, MBP-HIS-PL1, MBP-HIS-GFP, and SUMO-HIS-BIN2 proteins in *Escherichia coli*, cDNA of *POLAR, PL1*, or *GFP* were recombined into the pDEST-HIS-MBP vector. cDNA of *BIN2* was recombined into pET-SUMO-SH vector.

### Chemical treatments

For the single-cell sequencing experiment, 5 days post germination (dpg), Arabidopsis seedlings expressing *TMMpro:GUS-GFP* were immersed in liquid ½MS medium supplemented with 1 μM BL, 50 μM bikinin, or 0.1% (v/v) DMSO for 2 h. For the analysis of the effect of BL and bikinin on the abaxial cotyledon epidermis of Arabidopsis seedlings, seeds were directly germinated and grown for 3 days in liquid ½MS supplemented with the indicated concentration of BL or bikinin. For the BES1 phosphorylation assay upon BL or bikinin treatment, Arabidopsis seedlings of Col-0, *35Spro:gPOLAR*/Col-0, or *35Spro:gPL1*/Col-0 were germinated and grown on ½MS plates, for 5 days and then transferred to liquid ½MS for 2 h and supplemented with DMSO, 100 nM BL, or 50 μM bikinin for 1 h. For the root inhibition assay upon BL or bikinin treatment, Col-0, *35Spro:POLAR*/Col-0, or *35pro:gPL1*/Col-0 were germinated and grown on ½MS plates containing 0, 1, 5, or 10 nM BL for 5 days. For the nuclear and cytoplasmic BES1 fractionation assay, *35Spro:gPOLAR/BIN2pro:gBIN2-GFP/Col-0, 35Spro:gPL1/BIN2pro:gBIN2-GFP*/Col-0, and *BIN2pro:gBIN2-GFP*/Col-0 were grown on ½MS plate for 3 days and then transferred to liquid ½MS containing DMSO or 100 nM BL for 1 h.

### Single-cell sample preparation, library construction, and sequencing

Protoplasts were isolated as described previously^47^ with minor modifications. Briefly, the whole aerial tissue of the Arabidopsis seedlings was harvested and incubated in protoplasting solution [1.5% (w/v) cellulase Y-C, 0.15% (w/v) pectolyase Y-23, 0.4 M mannitol, 20 mM KCl, 10 mM CaCl_2_, 0.1% (w/v) bovine serum albumin (BSA), 20 mM MES, pH 5.7]. After 1 h of gentle shaking, cells were filtered through a 70-μm cell strainer and spun down at 100*g* for 6 min. Pelleted protoplasts were resuspended with washing solution [0.4 M mannitol, 20 mM KCl, 10 mM CaCl_2_, 0.1% (w/v) BSA, 20 mM MES, pH 5.7], and filtered through a 40-μm cell strainer. The protoplasts were then sorted on a FACSAria II (BD Biosciences) based on the GFP signal. Sorted cells were centrifuged at 400*g* at 4°C for 5 min and resuspended in washing solution to yield an estimated concentration of 1,000 cells/μl. Cellular suspensions were loaded on either a GemCode Single Cell 3′ Gel Bead and Library Kit (V2 chemistry, 10x Genomics) for the control experiment, or a Chromium Single Cell 3’ GEM, Library & Gel Bead Kit (V3 chemistry, 10× Genomics) for the treatment experiment according to the manufacturer’s instructions. Libraries were sequenced on a NovaSeq 6000 (Illumina) instrument following the 10x Genomics recommendations at the VIB Nucleomics Core facility (VIB, Leuven).

### ScRNA-seq dataset analyses

The raw sequencing data were demultiplexed with the 10x CellRanger (version 3.1.0) software ‘cellranger mkfastq’. The fastq files obtained after demultiplexing were used as the input for ‘cellranger count’, which aligns the reads to the Arabidopsis reference genome (Ensemble TAIR10.40) using STAR and collapses them to unique molecular identifier (UMI) counts. The result is a large digital expression matrix with cell barcodes as rows and gene identities as columns. Initial filtering in CellRanger recovered 13,796 cells for the control (*TMMpro:TMM-GFP*), 9,217 cells for the DMSO treatment, 8,384 cells for the BL treatment, and 8,198 cells for the bikinin treatment (*TMMpro:GFP*) samples, corresponding to 20,842 mean reads and 1,520 median genes, 37,828 mean reads and 2,132 median genes, 38,102 mean reads and 2,656 median genes, and 37,770 mean reads and 2,088 median genes per cell, respectively. To ensure that only high-quality cells were further analyzed, the filtered data provided by CellRanger were used as input for further filtering steps. All analyses were done in *R* (version > 3.6.0). Data were preprocessed by the scater package (version 1.10.1) following a recommended workflow^48^. Outlier cells were defined either as three median absolute deviations (MADs) away from the median value of the UMI numbers or numbers of expressed genes, or as cells containing more than 5% mitochondrial or 10% chloroplastic transcripts. After removal of the outliers, a final number was obtained of 8,180 cells for the control, 5,891 cells for the DMSO treatment, 7,460 cells for the BL treatment, and 6,803 cells for the bikinin treatment samples. Normalization of the raw counts, detection of highly variable genes, discovery of clusters, and creation of UMAP plots were done by means of the Seurat pipeline (version 4.0.3). Cells from the control dataset and the DMSO treatment were integrated into one object with the Harmony package^49^, in order to account for as many different cell types and states as possible. From this object, cells not belonging to the stomatal lineages (e.g., mesophyll and vascular cells) were removed. Differential expression analysis for marker gene identification per subpopulation was based on the nonparametric Wilcoxon rank sum test implemented within the Seurat pipeline. Clusters with the same cell annotation based on gene expression analysis were combined to generate a more comprehensible dataset. Similarly, the DMSO, BL, and bikinin datasets were integrated into one Seurat object with Harmony. This Seurat object was then converted to a SingleCellExperiment object and Slingshot^28^ was used to calculate a pseudotime lineage, using “UMAP” as reducedDim, “Pavement” as starting cluster, and “Mature GC” as end cluster. Cells not belonging to the trajectory were removed from the SingleCellExperiment object. The generalized additive models were calculated with the fitGAM command from the tradeSeq package^29^ with the different samples as conditions and with six knots, which were determined by the evaluateK function. Expression patterns behaving differently between the treatments were determined with the conditionTest function of tradeSeq and visualized with the plotSmoothers function.

### BES1 dephosphorylation assay and quantification

Arabidopsis seedlings of Col-0, *35Spro:gPOLAR*/Col-0, or *35Spro:gPL1*/Col-0 were germinated and grown on ½MS plates for 5 days and then transferred to liquid ½MS for 2 h and supplemented with 100 nM BL or 50 μM bikinin for 1 h. After treatment, seedlings were harvested and frozen in liquid nitrogen. For the BES1 dephosphorylation analysis, total proteins were extracted with buffer containing 25 mM Tris-HCl, pH 7.5, 150 mM NaCl, 1% (w/v) sodium dodecyl sulfate (SDS), 10 mM dithiothreitol (DTT), and EDTA-free protease inhibitor mixture complete (Roche Diagnostics). For blocking and antibody dilutions, 5% (w/v) skim milk powder in Tris-buffered saline containing 0.2% (v/v) TWEEN20 was used. Proteins were resolved by 10% SDS-polyacrylamine gel electrophoresis (PAGE) and detected by western blot with polyclonal anti-BES1^3^ and anti-tubulin (Sigma-Aldrich). Dephosphorylated BES1, phosphorylated BES1, and tubulin proteins were quantified based on the signal intensity with ImageJ (https://imagej.nih.gov/ij/).

### Quantitative real-time PCR (qRT-PCR)

RNA was extracted from 100 mg of seedlings at 3 dpg by means of the RNeasy mini kit (Qiagen) according to the manufacturer’s instructions. cDNA was generated with the qScript cDNA SuperMix (Quantabio). For the qRT-PCR; a LightCycler® 480 machine (Roche Diagnostics) was used with SYBR green I qPCR master mix (Roche Diagnostics). For the validation of gene expression identified in the single-cell RNA sequencing analysis, RNA was extracted from 20,000 to 50,000 GFP-positive cells sorted from the protoplasts of Arabidopsis cotyledons expressing *TMMpro:GUS-GFP*. Cotyledons were harvested at 5 dpg after treatment with 1 μM BL, 50 μM bikinin, or 0.1% (v/v) DMSO for 2 h. Primers are listed in Supplementary Table 2.

### Quantitative analysis of epidermal phenotypes

For epidermal cell analysis of developing cotyledons of Arabidopsis seedlings at 3 dpg, cotyledons_were stained by propidium iodide (50 μg/mL) and imaged with a SP8 confocal microscope (Leica). Epidermal cell surface areas in a 0.125-mm^2^ region from the central part of a cotyledon were measured in ImageJ with free-hand selection tool. Cells smaller than 200 μm^2^ were defined as small nonstomatal cells, whereas cells larger than 200 μm^2^ were defined as pavement cells. Small nonstomatal cell density (number of small nonstomatal cells/mm^2^), pavement cell density (number of pavement cells/mm^2^), total cell density (total number of cells/mm^2^), and stomata index (number of stomata/total number of cells × 100) were calculated.

### Microscopy, image acquisition, and image analysis

The abaxial side of cotyledons of Arabidopsis seedlings at 3 dpg or infiltrated *N. benthamiana* leaves were analyzed with a SP8 confocal microscope (Leica). The images were taken at 405 nm, 488 nm, and 559 nm laser excitation and 425-460 nm, 500-530 nm, and 570-670 nm long-pass emission for TagBFP2, eGFP, and mCherry/propidium iodide/FM 4-64, respectively. The gating system was applied for autofluorescence removal. The *z* stack images were taken with an interval of 4 μm. For cotyledon epidermis staining, N-(3-triethylammoniumpropyl)-4-(6-(4-(diethylamino) phenyl) hexatrienyl) pyridinium dibromide (FM 4-64; 50 μM) (Invitrogen) or propidium iodide (PI) (50 μg/ml) (Sigma-Aldrich) was used as a plasma membrane marker.

For time-lapse imaging, cotyledons of Arabidopsis seedlings at 2 dpg were mounted in a chamber filled with ½MS agar. The *z* stack images were captured at 40-min intervals during the time indicated. The movies were made at the speed of three frames per second (f.p.s). For quantification of the nucleus BES1-GFP intensity, the Fiji image analysis software (http://fiji.sc) was used. For the measurement of fluorescence intensity changes upon BL or bikinin treatment, the abaxial side of cotyledons at 5 dpg was analyzed. All the fluorescent nuclei were outlined with the ‘Analyze particles’ function of the Fiji software. The mean gray value of the each selected area was then measured.

### Co-immunoprecipitation (Co-IP) assay

The plant materials were ground into powder with liquid nitrogen and resuspended in extraction buffer [50 mM Tris-HCl, pH 7.5, 50 mM NaCl, 300 mM sucrose, 1% (v/v) Triton X-100, 1× protease inhibitor, and 0.2 mM phenylmethylsulfonyl fluoride (PMSF)]. After centrifugation at 15,000 □ *g* for 10 min, the supernatants were incubated with GFP trap magnetic agarose for 1 h. Beads were washed four times with extraction buffer containing 0.2% (v/v) Triton X-100 and eluted with 2× SDS sample buffer [24 mM Tris-HCl, pH 6.8, 10% (v/v) glycerol, 0.8% (w/v) SDS, and 2% (v/v) 2-mercaptoethanol].

### Purification of bacterially produced proteins

The recombinant plasmids of *pDEST-HIS-MBP-POLAR, pDEST-HIS-MBP-PL1, pDEST-HIS-MBP-GFP*, and *pET-SUMO-SH-BIN2* were transformed into *E. coli* BL21 Rosetta (DE3) cells. MBP-HIS-POLAR, MBP-HIS-PL1, and MBP-HIS-GFP proteins were purified with amylose resin (NEB) and SUMO-HIS-BIN2 protein was purified with Ni-NTA Agarose (Qiagen).

### *In vitro* kinase assay

MBP bead-bound MBP-HIS-GFP, MBP-HIS-POLAR, and MBP-HIS-PL1 proteins were incubated with SUMO-HIS-BIN2 in kinase assay buffer (50 mM Tris-HCl, pH 7.5, 100 mM NaCl, 10 mM MgCl2, and 1 mM DTT), 100 μM cold ATP, and 5 μCi (γ-^32^P) ATP for 2 h at 30□. After the reaction, each supernatant fraction of SUMO-HIS-BIN2 was collected and separated by SDS-PAGE.

### Nuclear and cytoplasmic fractionation

Subcellular fractionation was carried out as described previously^50^. Arabidopsis seedlings were ground in liquid nitrogen and extracted with lysis buffer [20 mM HEPES, pH 7.5, 40 mM KCl, 10% (v/v) glycerol, 1 mM EDTA, 10 mM MgCl_2_, and 1% (v/v) Triton X-100]. Each extract was filtered and centrifuged at 5000*g* for 2 min. The supernatants were collected as cytoplasmic fractions and the pellets were washed several times with the buffer. Clear nuclear pellets were resuspended with the same volume as that of the cytosolic fraction. Nuclear and cytosolic fractions were mixed with SDS sample buffer and separated by SDS-PAGE. Anti-H3 and anti-PEPC were applied to detect nuclear and cytoplasmic marker, respectively.

### Statistical analysis

All statistical analyses in this paper were performed in GraphPad Prism 9 software except for the box plots. All the box plots were generated by BoxPlotR^51^ and *P* values were calculated by one-way ANOVA. Comparisons of one genotype were done by a single factor ANOVA with Dunnett’s multiple comparisons test. Comparisons of more than two genotypes were done by two-way ANOVA with Tukey’s multiple comparisons test. Details of statistical analyses are provided in the figure legends.

## Supporting information

Data Set1

Data Set2

## Data availability

All data are available in the paper or the Supplementary Information. The scRNA-seq datasets are available at the NCBI Gene Expression Omnibus repository under accession number GSE193451.

## Acknowledgments

We thank Y. Yin (Iowa State University) for the BES1 antibody, A. Vatén (University of Helsinki), X. Wang (Henan University), C. Fenoll (University of Castilla-La Mancha), T. Kakimoto (Osaka University), D. Bergmann (Stanford University), K. Torri (University of Texas at Austin) and L. De Veylder (VIB-UGent) for sharing published materials, N. Vukašinović for useful discussion and Martine De Cock for help in preparing the manuscript. The *CYCD7;1pro:GFP-GUS* line was kindly provided by L. De Veylder (VIB-UGent). This work is supported by Research Foundation-Flanders (project G003720N to E. R. and a postdoctoral fellowship 1222221N to E.-J. K.), Chinese Scholarship Council (predoctoral fellowships to C. Z. and B. G. and a visiting scientist fellowship to K.W.), Belgian Science Policy (postdoctoral fellowship M. T. and K. W.), Ghent University “Bijzondere Onderzoekfonds” (BOF/CHN/010 to C.Z. and BOF18/DPO/151 to J.R.W.), and European Research Council (ERC Starting Grant TORPEDO, no. 714055 to B.D.R.).

## Author contributions

A. H., C.Z., K. W., B.D.R and E.R. conceived the project. E.-J.K., C.Z., B.G., A.H., M.T., C. S.--V., and I.V. performed experiments and analysed data. B.G., T.E., J.R.W., N.V., Y.S., and B.D.R performed the scRNA-seq and data analysis. Y.Z. and K.W. supervised data analysis. E.-J.K., C.Z., B.G., T.E., B.D.R. and E.R. wrote the manuscript and all authors revised it.

---

**Extended Data Fig. 1.**
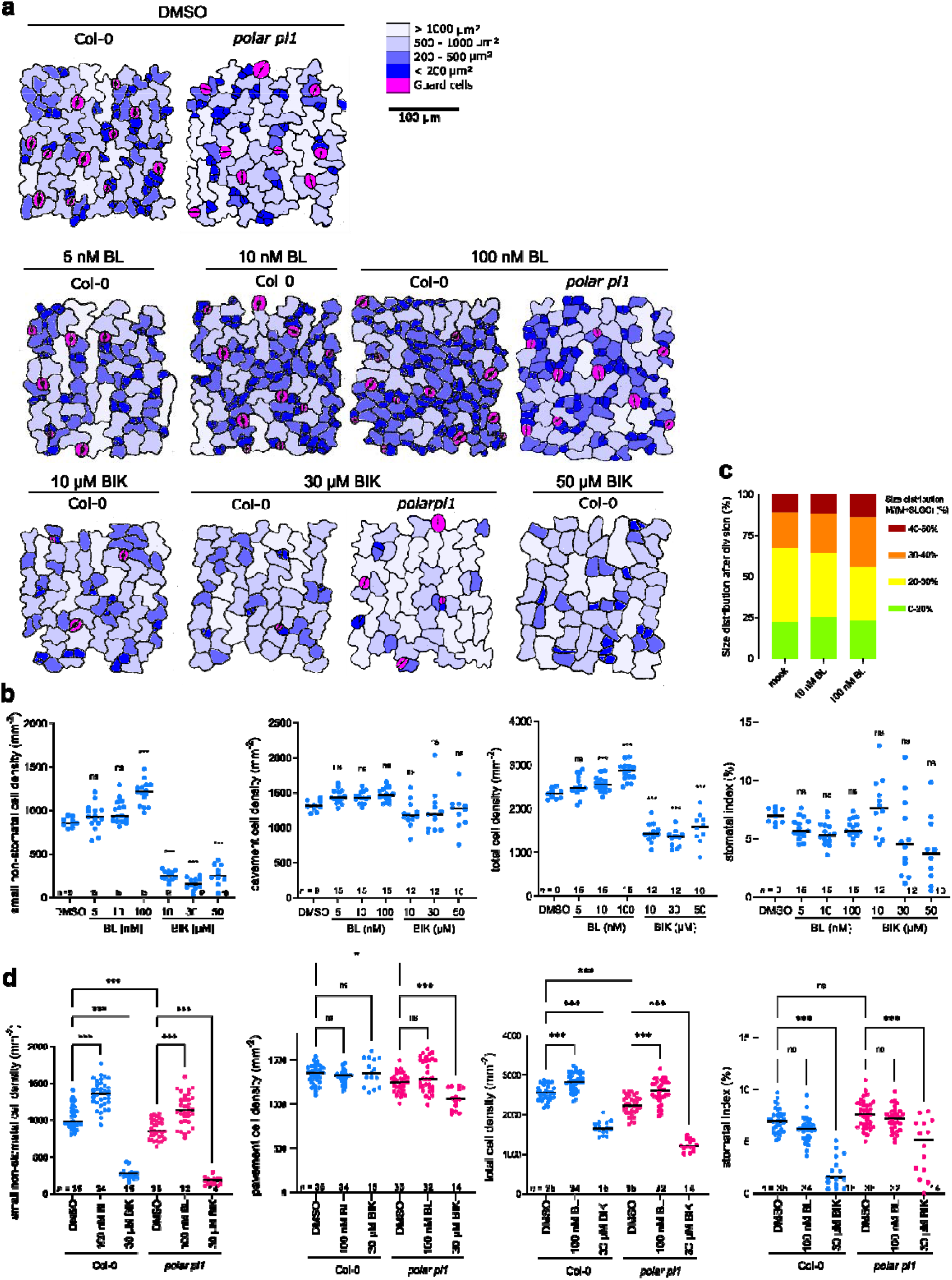
BRs and bikinin inversely affect stomatal development in Arabidopsis. **a,** Abaxial cotyledon epidermis of Arabidopsis wild type Col-0 and *polar pl1* double mutant. Seeds were germinated and grown for 3 days in liquid media supplemented with brassinolide (BL) and bikinin (BIK). DMSO was used as a mock control. Cell size distribution is presented with a colour scale. Scale bar, 100 μm. **b,** Quantification of epidermal phenotypes of the wild type in **a**. Graphs represent small (<200 μm^2^) non-stomatal cell density, pavement cell (≧200 μm^2^) density, total cell density, and stomatal index. All individual data points are shown and black horizontal bars represent the means. Asterisks highlight significant differences from the DMSO treatment by one-way ANOVA and Dunnett’s multiple comparisons tests. \*\*\**P*< 0.001; ns, not significant; *n*, number of cotyledons analysed. The data shown are from one experiment. **c,** Quantification of size distribution between meristemoids (Ms) and stomatal lineage ground cells (SLGCs) after cell division of the wild type Col-0. The area of each cell was measured and classified into four groups as indicated by the size distribution between meristemoids and SLGCs after cell division (%). From each genotype, 100 meristemoids and 100 SLGCs were analysed. **d,** Quantification of epidermal phenotypes of the *polar pl1* double mutant in **a**. Two-way ANOVA and Tukey’s multiple comparisons test with a single pooled variance was performed. \*\*\**P*< 0.001, \**P*< 0.05; ns, not significant; *n*, number of cotyledons analysed. Data from two independent experiments are shown.

**Extended Data Fig. 2.**
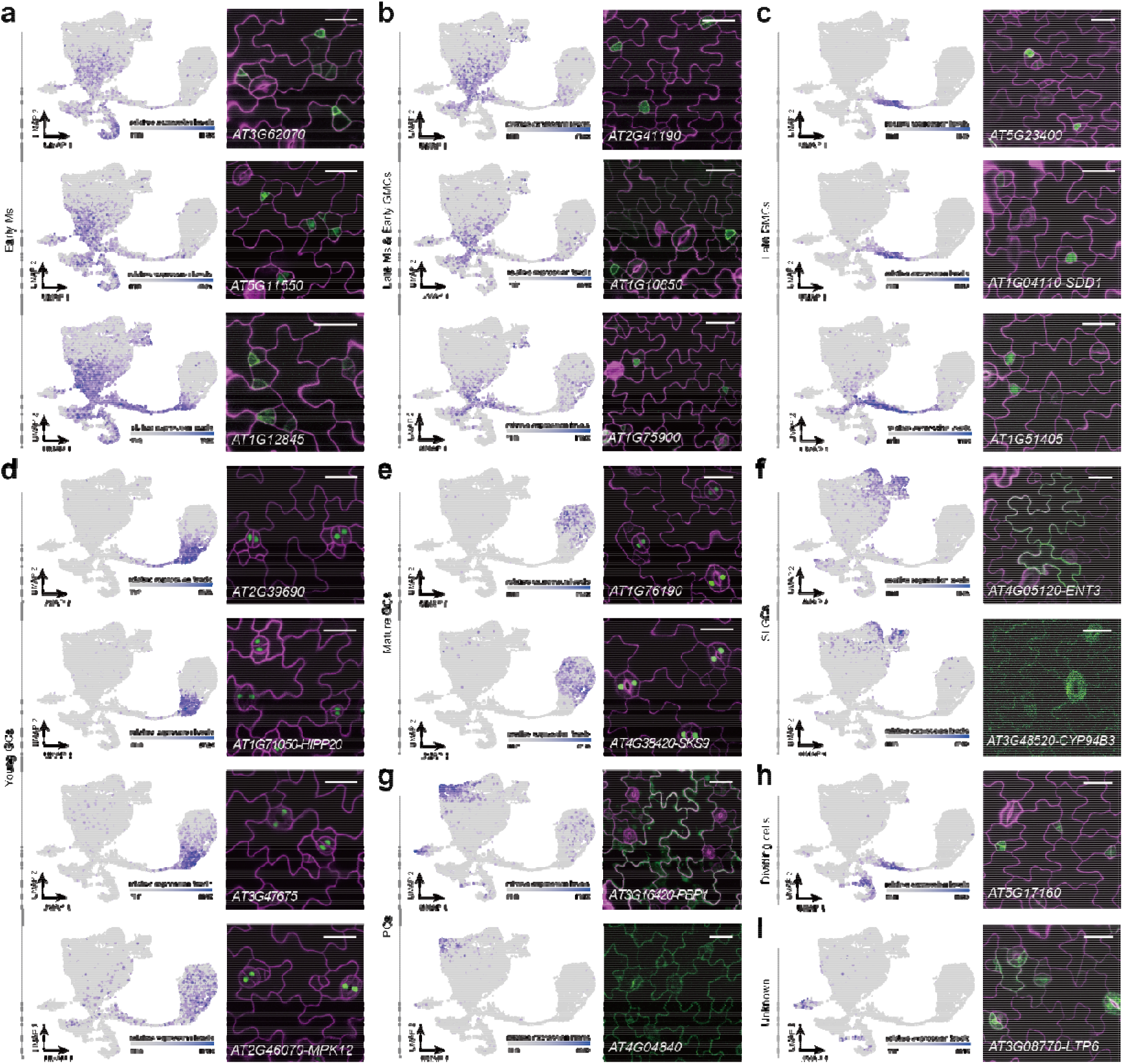
Validation of the control cluster annotation dataset. **a-i,** Validation of the cell cluster annotation, including feature plots and reporter line expression patterns of differentially expressed genes (DEGs) for each of the ten clusters. Constructs containing the promoter region upstream of the transcriptional start for each gene driving the expression of the full-length genomic coding region fused to GFP were used except for *AT2G39690, AT1G71050, AT3G47675, AT2G46070, AT1G76190*, and *AT4G38420*, for which promoter-GFP fusions were generated. The abaxial cotyledon epidermis was imaged at 5 days post germination. Cell outlines (magenta) were marked with *ML1pro:mCherry-RCI2A* for *AT5G23400, AT2G39690, AT1G71050, AT3G47675, AT2G46070, AT1G76190, AT4G38420*, and *AT3G08770* and with propidium iodide in the other images. Scale bars, 25 μm. M, meristemoid; GMC, guard mother cell; SLGC, stomatal lineage ground cell; GC, guard cell; PC, pavement cell.

**Extended Data Fig. 3.**
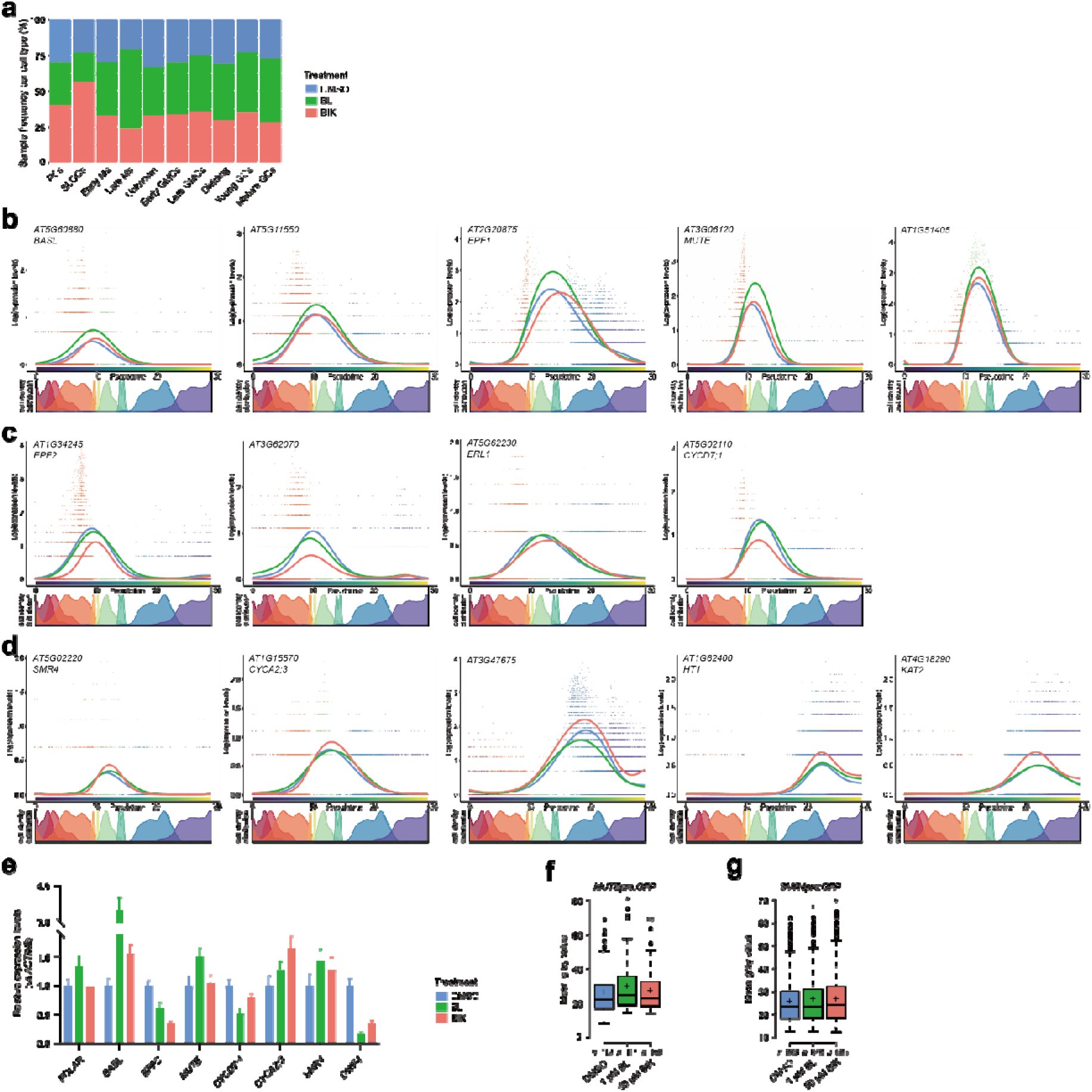
Gene expression of stomatal markers across developmental trajectories and treatments. **a,** Sample contribution of the merged dataset across each cell cluster. **b-d,** Expression dynamics of stomatal genes that are upregulated by brassinolide (BL) (**b**), downregulated by bikinin (BIK) (**c**), and upregulated by BIK (**d**) along a pseudotime. **e,** Quantitative real-time PCR analysis of stomatal genes in response to BL and bikinin. Total RNA was extracted from 20,000-50,000 GFP-positive cells sorted from protoplasts isolated from *TMMpro:GFP* expressing Arabidopsis plants treated with 1 μM BL, 50 μM bikinin, and DMSO (mock) for 2 h in liquid medium. Transcript levels were normalized to *ACTIN2* (AT3G18780). *DWARF4* (*DWF4*, AT3G50660) was used as a positive control for the treatments. Error bars represent SD. One biological replicate with three technical replicates were analysed. **f,g,** Fluorescence intensity of *MUTEpro:GFP* (**f**) and *SMR4pro:GFP* (**g**) in response to BL and bikinin. Arabidopsis plants at 5 days post germination were imaged after 2 h of treatment with 1 μM BL, 50 μM bikinin, and DMSO (mock) in liquid medium. The mean grey value of all the fluorescent nuclei was measured with ImageJ. Center lines show the medians. Crosses represent sample means. Box limits indicate the 25th and 75th percentiles as determined by R software. Whiskers extend 1.5 times the interquartile range from the 25th and 75th percentiles. Outliers are represented by dots. Asterisks highlight significant differences from the DMSO treatment determined with a single factor ANOVA. \**P*< 0.05. ns, no significant difference. *n*, number of fluorescent nuclei measured.

**Extended Data Fig. 4.**
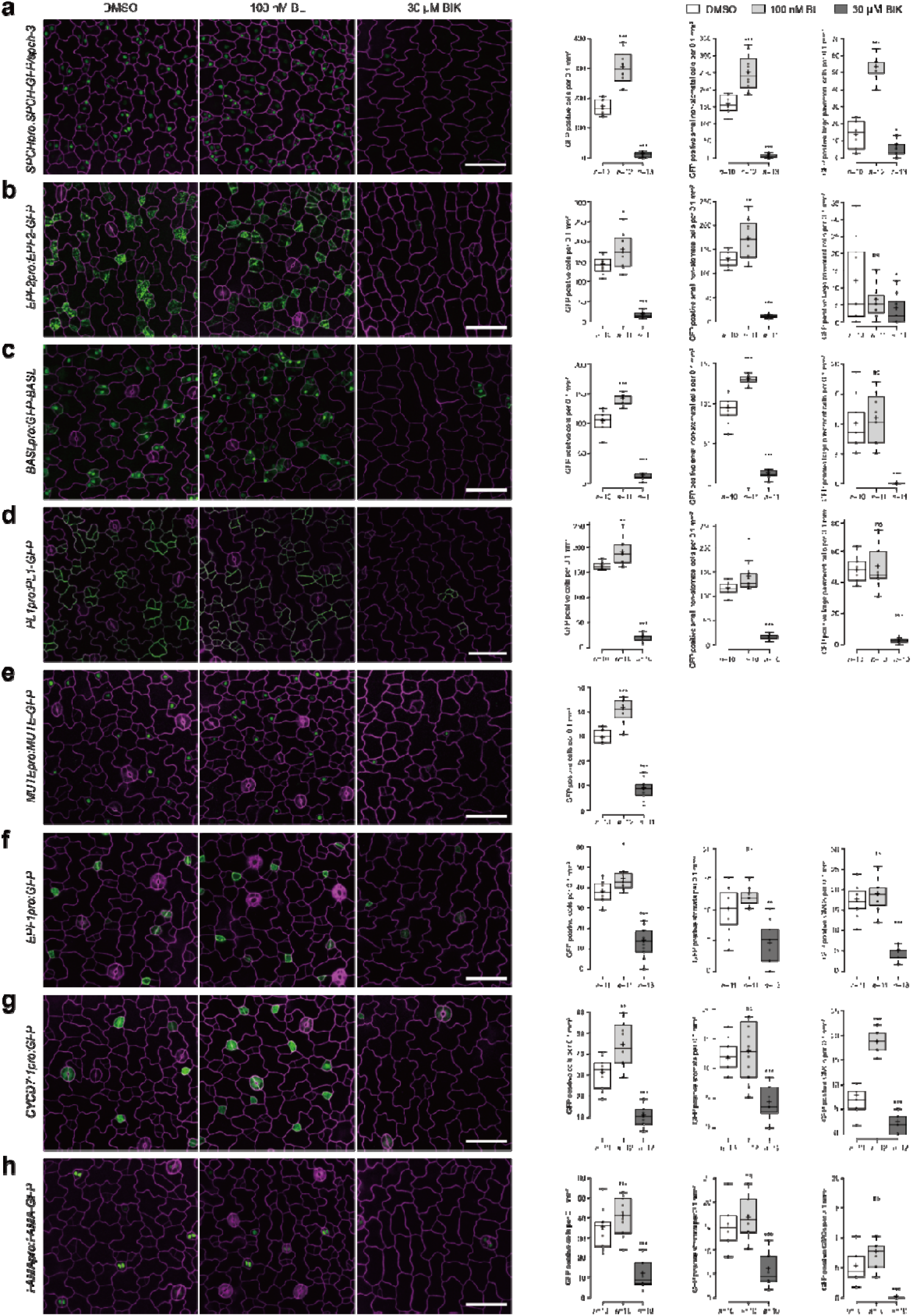
The expression of stomatal markers in response to BL and bikinin treatment. **a-h,** The expression of SPCH (**a**), EPF2 (**b**), BASL (**c**), PL1 (**d**), MUTE (**e**), EPF1 (**f**), CYCD7;1 (**g**), and FAMA (**h**) after treatment with brassinolide (BL), bikinin (BIK), and DMSO (mock). Seeds were germinated and grown in liquid medium containing 100 nM BL, 30 μM bikinin or DMSO for 3 days. Cell outlines (magenta) were marked with *UBQ10pro:PIP2A-mCherry* for BASL (**c**) and with propidium iodide in the other panels. Scale bars, 50 μm. GFP-positive cells were calculated as small (<200 μm^2^) non-stomatal cells and large pavement cells (≧200 μm^2^) per 0.1 mm^2^ for SPCH (**a**), EPF2 (**b**), BASL (**c**), PL1 (**d**). Data points are plotted as dots. Center lines show the medians. Crosses represent sample means. Box limits indicate the 25th and 75th percentiles as determined by R software. Whiskers extend 1.5 times the interquartile range from the 25th and 75th percentiles. Asterisks highlight significant differences from the DMSO treatment determined with a single factor ANOVA. \*\*\**P*< 0.001, \*\**P*< 0.01, \**P*< 0.05; ns, no significant difference. *n*, number of cotyledons analysed.

**Extended Data Fig. 5.**
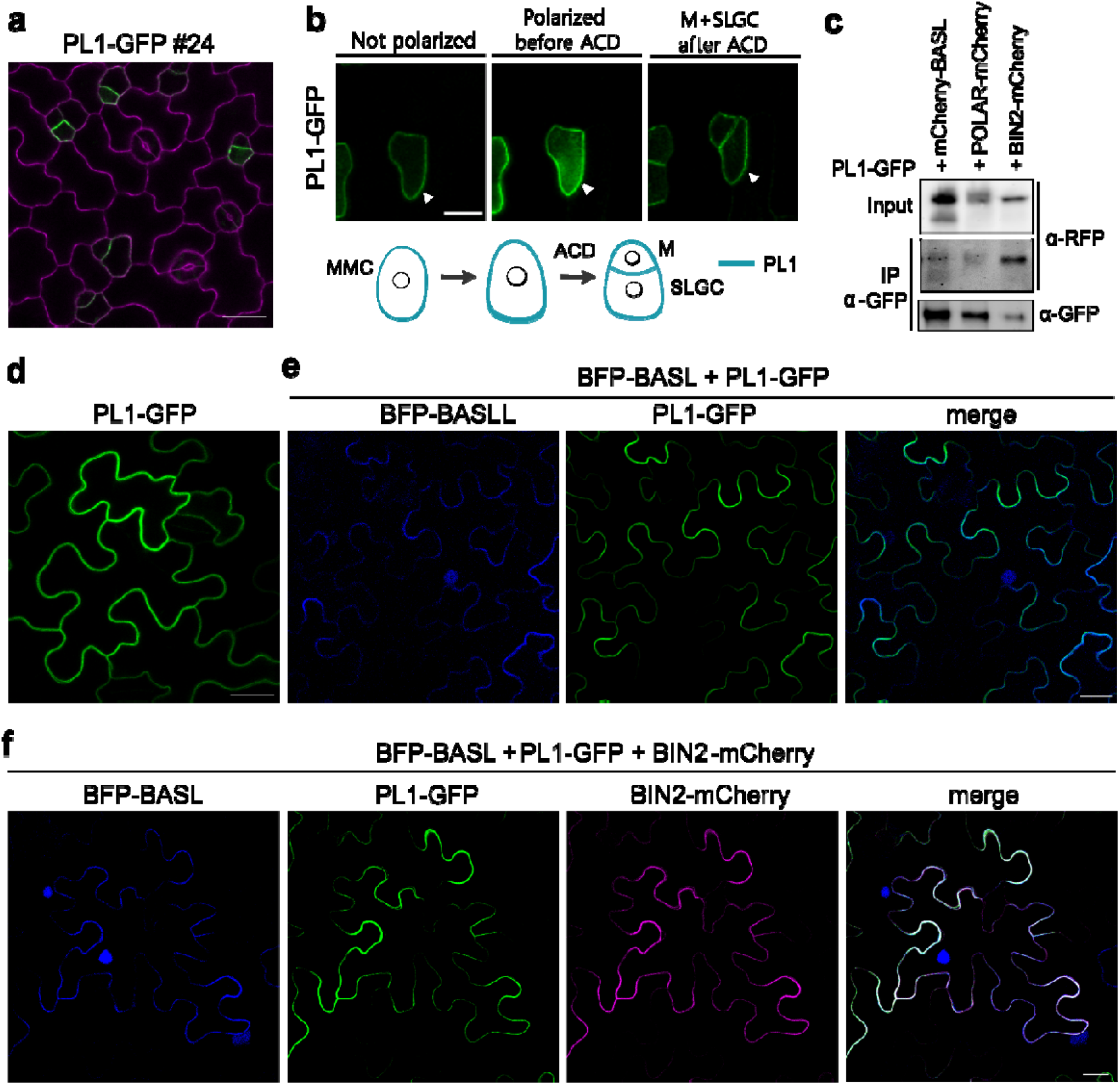
POLAR-LIKE1 (PL1) functions together with POLAR. **a,** Expression pattern of PL1 in stomatal lineage cells. The abaxial cotyledon epidermis of Arabidopsis plants expressing PL1-GFP [*PL1pro:gPL1-GFP*/Col-0 (line #24)] at 3 days post germination (dpg) was examined by confocal microscopy. Cell outlines (magenta) were marked with propidium iodide. Scale bar, 20 μm. **b,** Time-lapse imaging of PL1-GFP. The abaxial cotyledon epidermis of *PL1pro:gPL1-GFP*/Col-0 (line #24) at 2 dpg were observed by time-lapse imaging. The images were taken at an interval of 1 h. Arrows indicate protein polarization. Scale bar, 10 μm. The cartoon shows the localization of PL1-GFP during asymmetric cell division (ACD) in the stomatal lineage. Blue, PL1; MMC, meristemoid mother cell; M, meristemoid; SLGC, stomatal lineage ground cell. **c,** mCherry-BASL, POLAR-mCherry, and BIN2-mCherry co-immunoprecitated with PL1-GFP when transiently co-expressed in tobacco leaf epidermis. Protein extracts from tobacco leaf expressing PL1-GFP together with mCherry-BASL, POLAR-mCherry, or BIN2-mCherry, were immunoprecipitated by α-GFP and immunoblots were detected by α-GFP and α-RFP antibodies. IP, immunoprecipitation. **d,** Transient expression in tobacco leaf epidermis of PL1-GFP. **e,** Transient co-expression of PL1-GFP with BFP-BASL strongly polarizing PL1 in tobacco leaf epidermis. **f,** Transient co-expression of BIN2-mCherry with PL1-GFP and BFP–BASL strongly polarizing BIN2 and PL1 in leaf epidermis of tobacco. Scale bars, 20 μm in **d-f**.

**Extended Data Fig. 6.**
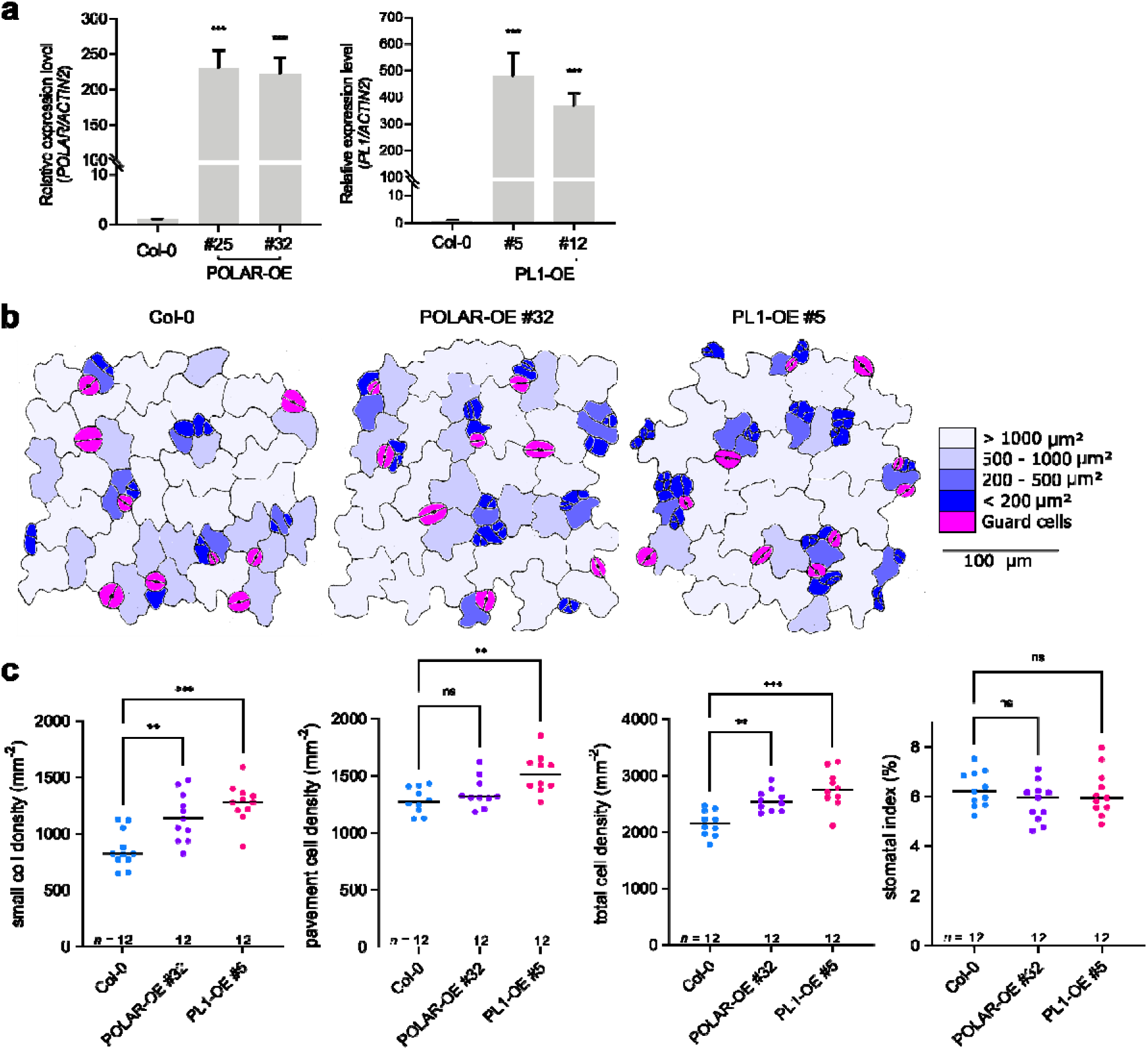
Stomatal phenotype of *POLAR* and *PL1* overexpressing plants. **a,** Transcript levels of *POLAR* and *PL1* analysed with qRT-PCR in Arabidopsis wild type (Col-0) and plants overexpressing *POLAR (35Spro:gPOLAR*, POLAR-OE, lines #25 and #32) and *PL1* (*35Spro:gPL1*, PL1-OE, lines #5 and #12). Total RNA was extracted from seedlings 3 days post germination (dpg). Transcript levels were normalized to *ACTIN2* (AT3G18780). Two biological replicates with three technical replicates were analysed. Error bars represent SD. **b,** Representation of the abaxial cotyledon epidermis of Arabidopsis Col-0, POLAR-OE [*35Spro:gPOLAR*/Col-0 (line #32)], and PL1-OE [*35Spro:gPL1*/Col-0 (line #5)] plants at 3 dpg. Cell size distribution is presented as a colour scale. Scale bar, 100 μm. **c,** Quantification of epidermal phenotypes in **b**. Graphs represent the small (<200 μm^2^) nonstomatal cell density, pavement cell (≧200 μm^2^) density, total cell density, and stomatal index. All individual data points are shown and black horizontal bars represent the means. The significant differences between the transgenic lines and Col-0 control were determined by one-way ANOVA and Dunnett’s multiple comparisons tests. \*\*\**P*< 0.001, \*\**P*< 0.01; ns, not significant. *n*, number of biologically independent cotyledons.

**Extended Data Fig. 7.**
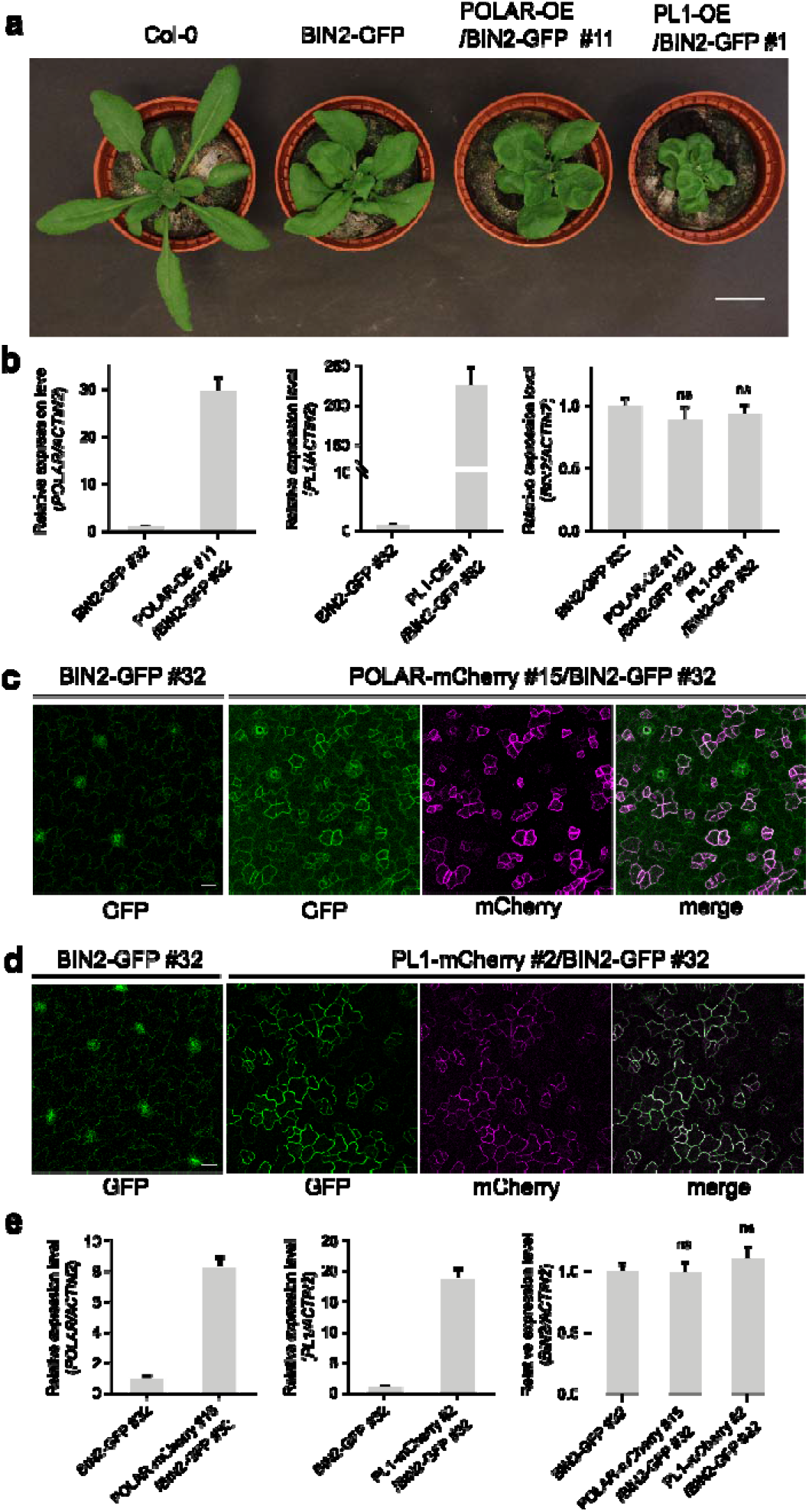
Characterization of the transgenic lines overexpressing either POLAR or PL1 together with BIN2-GFP. **a,** Four-week-old Arabidopsis wild type (Col-0), BIN2-GFP [*BIN2pro:gBIN2-GFP*/Col-0 (line #32.7)], POLAR-OE/BIN2-GFP [*35Spro:gPOLAR/BIN2pro:gBIN2-GFP* (line #32.7)/Col-0 (line #11)], and PL1-OE/BIN2-GFP [*35Spro:gPL1/BIN2pro:gBIN2-GFP* (line #32.7)/Col-0 (line #1)] plants. Scale bar, 2 cm. **b,** Transcript levels of *POLAR, PL1*, and *BIN2* in plants shown in **a** and analysed by qRT-PCR. Two biological replicates with three technical replicates were analysed. Error bars represent SD. ns, not significant. **c,d,** Expression patterns of BIN2-GFP [*BIN2pro:gBIN2-GFP*/Col-0 (line #32.7)], and BIN2-GFP combined with either POLAR-mCherry [*POLARpro:gPOLAR-mCherry/BIN2pro:gBIN2-GFP* (line #32.7)]/Col-0 (line #15)] or PL1-mCherry [*PL1pro:gPL1-mCherry/BIN2pro:gBIN2-GFP*(line #32.7)]/Col-0 (line #2]. The abaxial cotyledon epidermis of Arabidopsis seedlings at 3 days post germination (dpg) was observed by confocal microscopy. Scale bars, 20 μm. **e,** Transcript levels of *POLAR, PL1*, and *BIN2* in plants described in **c** and **d** analysed by qRT-PCR. In **b** and **e** total RNA was extracted from seedlings at 3 dpg. Transcript levels were normalized to *ACTIN2* (AT3G18780). Two biological replicates with three technical replicates were analysed. Error bars represent SD. ns, not significant.

**Extended Data Fig. 8.**
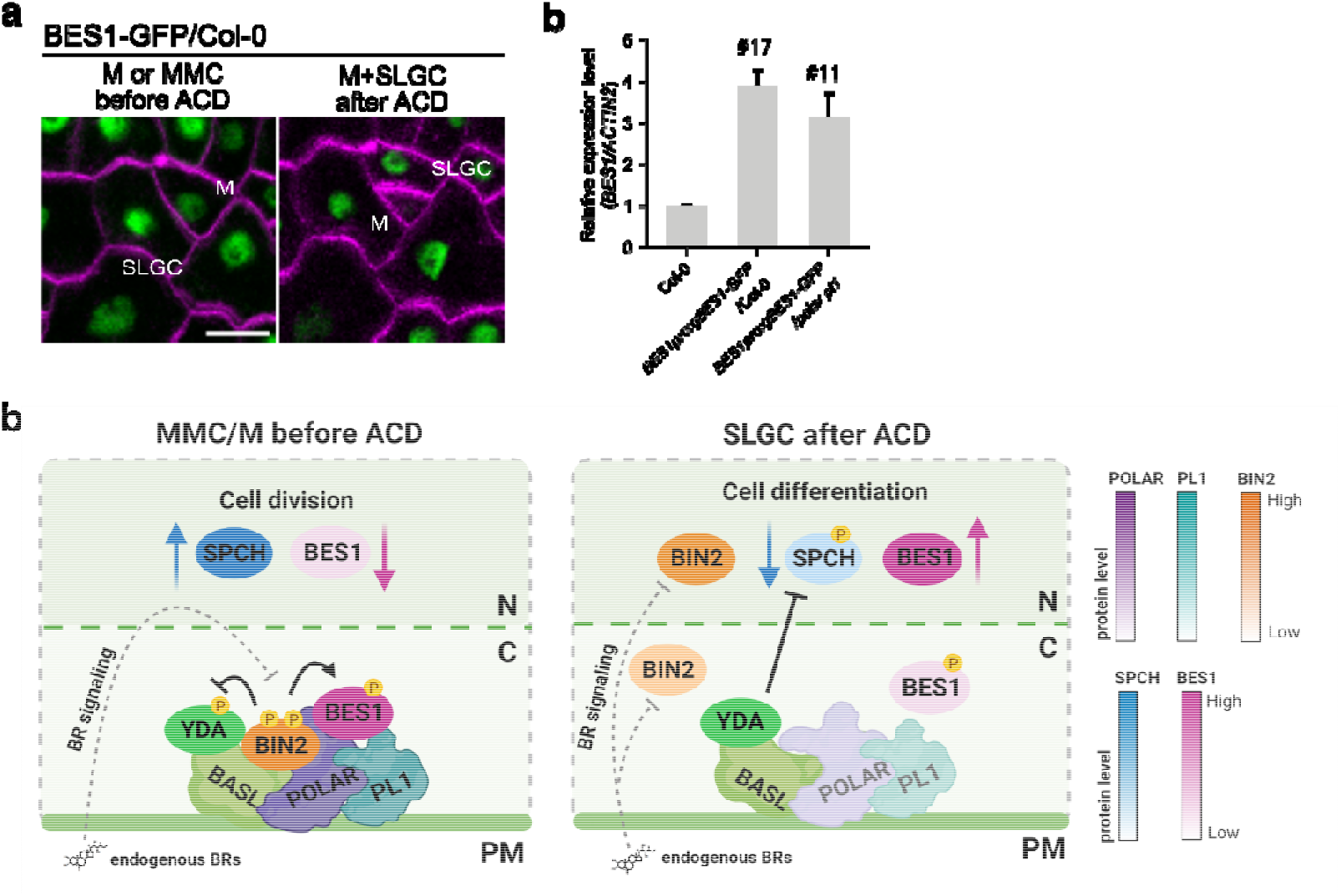
Regulation of BR signalling in the stomatal lineage. **a,** Expression pattern of BES1-GFP [*BES1pro:gBES1-GFP*/Col-0 (Line #17)] in wild type Arabidopsis before and after stomatal asymmetric cell division (ACD). The abaxial cotyledon epidermis at 2 days post germination (dpg) was examined by time-lapse imaging. The experiment was repeated independently three times. Images were taken at an interval of 1 h Scale bar, 10 μm. **b,** Transcript levels of *BES1* in Col-0, *BES1pro:gBES1-GFP*/Col-0 (line #17), and *BES1pro:gBES1-GFP/polar pl1* (line #11) analysed by qRT-PCR. Total RNA was extracted from seedlings at 3 dpg. Transcript levels were normalized to the *ACTIN2* (AT3G18780) gene. Two biological replicates with three technical replicates were analysed. Error bars represent SD. **c,** Model for BR signalling regulation in the stomatal lineage. In the meristemoid mother cell (MMC) or meristemoid (M), in which POLAR and PL1 are highly expressed before the ACD, BIN2 is polarized and highly enriched in the plasma membrane (PM) together with BASL, POLAR, and PL1. In this complex, BIN2 remains active, insensitive to BR-mediated inhibition and blocks the MAPK signalling module. As a result, the high activity of SPCH in the nucleus promotes ACD. BES1 is associated with POLAR and PL1 in the PM, remains phosphorylated by BIN2 and thus, BR responses are suppressed. After ACD, the expression of POLAR and PL1 in stomatal lineage ground cells (SLGCs) is reduced leading to BIN2 dissociation from the PM and, as a consequence, BIN2 is accessible for BR-mediated inactivation. Hence, the MAPK signalling is activated causing a strong suppression of SPCH activity, which restricts ACD. Dephosphorylated BES1 is then translocated into the nucleus to activate BR responses followed by pavement cell differentiation. Dotted rectangles represent the part of MMC/M (left) and SLGC (right). N, nuclear. C, cytoplasm. The BL molecules represent endogenous BRs. The protein levels of POLAR, PL1, BIN2, SPCH, and BES1 are depicted with colour intensity. The figure is created with BioRender (BioRender.com).

