## Supplementary material for "Cell type-specific attenuation of brassinosteroid signalling drives stomatal asymmetric cell division": SI

### Supplementary information

### Supplementary Table 1. The constructs generated for the validation of cell cluster identities

| Cluster | Gene identifier | Gene description | Destination construct | Base vector |
|---|---|---|---|---|
| Early Ms | AT3G62070 | Unknown | *AT3G62070pro:GFP-gAT3G62070* | pFASTRK-AG |
| Early Ms | AT5G11550 | ARM repeat superfamily protein | *AT5G11550pro:GFP-gAT5G11550* | pB7m34GW |
| Early Ms | AT1G12845 | Transmembrane protein | *AT1G12845pro:gAT1G12845-GFP* | pFASTRK-AG |
| Late Ms/early GMCs | AT2G41190 | Transmembrane amino acid transporter family protein | *AT2G41190pro:gAT2G41190-GFP* | pB7m34GW |
| Late Ms/early GMCs | AT1G75900 | GDSL-motif esterase/acyltransferase/lipase | *AT1G75900pro:GFP-gAT1G75900* | pB7m34GW |
| Late Ms/early GMCs | AT1G10850 | Leucine-rich repeat protein kinase family protein | *AT1G10850pro:gAT1G10850-GFP* | pFASTRK-AG |
| Late GMCs | AT1G04110 | SDD1, encodes a protein similar to serine proteases | *AT1G04110pro:GFP-gAT1G04110* | pB7m34GW |
| Late GMCs | AT5G23400 | Leucine-rich repeat family protein | *AT5G23400pro:GFP-gAT5G23400* | pFASTRK-AG |
| Late GMCs | AT1G51405 | Myosin-like protein | *AT1G51405pro:GFP-gAT1G51405* | pB7m34GW |
| Young GCs | AT2G39690 | Ternary complex factor MIP1 leucine-zipper protein | *AT2G39690pro:nlsGFP* | pGGB-AG |
| Young GCs | AT1G71050 | HIPP20, heavy metal transport/detoxification superfamily protein | *AT1G71050pro:nlsGFP* | pGGB-AG |
| Young GCs | AT3G47675 | Ternary complex factor MIP1 leucine-zipper protein | *AT3G47675pro:nlsGFP* | pGGB-AG |
| Young GCs | AT2G46070 | MPK12, encodes a MAP kinase protein | *AT2G46070pro:nlsGFP* | pGGB-AG |
| Mature GCs | AT1G76190 | SAUR56, SAUR-like auxin-responsive protein family | *AT1G76190pro:nlsGFP* | pGGB-AG |
| Mature GCs | AT4G38420 | SKS9, SKU5 similar 9 | *AT4G38420pro:nlsGFP* | pGGB-AG |
| SLGCs | AT4G05120 | ENT3, encodes an equilibrative nucleoside transporter | *AT4G05120pro:GFP-gAT4G05120* | pFASTRK-AG |
| SLGCs | AT3G48520 | CYP94B3, jasmonoyl-isoleucine-12-hydroxylase | *AT3G48520pro:GFP-gAT3G48520* | pFASTRK-AG |
| Pavement cells | AT4G04840 | MSRB6, methionine sulfoxide reductase B6 | *AT4G04840pro:gAT4G04840-GFP* | pB7m34GW |
| Pavement cells | AT3G16420 | PBP1, assists the PYK10 (β-glucosidase complex) in its activity | *AT3G16420pro:gAT3G16420-GFP* | pFASTRK-AG |
| Unknown | AT3G08770 | LTP6, predicted to encode a PR (pathogenesis-related) protein, belongs to the lipid transfer protein (PR-14) family | *AT3G08770pro:GFP-gAT3G08770* | pB7m34GW |
| Dividing cells | AT5G17160 | aspartic/glutamic acid-rich protein | *AT5G17160pro:gAT5G17160-GFP* | pFASTRK-AG |

Ms, meristemoids; GMCs, guard mother cells; GCs, ground cells; SLGCs, stomatal lineage ground cells

### Supplementary Table 2. Primers used in this study

| Name | Sequence |
|---|---|
| **Oligos for cloning** | |
| pPOLARlike1_AttB4_Fw | GGGGACAACTTTGTATAGAAAAGTTGGGGAAGGCGCATATGCGGTTCT |
| pPOLARlike1-ORF2-AttB1R_Rv | GGGGACTGCTTTTTTGTACAAACTTGTCGGAGGATTCAAAATTCGATC |
| POLARlike1-ORF2-AttB1_Fw | GGGGACAAGTTTGTACAAAAAAGCAGGCTTTATGGATGACGGTGTAGGAGTG |
| POLARlike1_AttB2_Stop_Rv | GGGGACCACTTTGTACAAGAAAGCTGGGTATCACCTTGAGAGAGGGTAAG |
| POLARlike1_AttB2_no Stop_Rv | GGGGACCACTTTGTACAAGAAAGCTGGGTACCTTGAGAGAGGGTAAGGAT |
| pBES1-gBES1_attB4F | GGGGACAACTTTGTATAGAAAAGTTGGGtatattagtatcactattct |
| pBES1-gBES1_attB1R | GGGGACTGCTTTTTTGTACAAACTTGTACTATGAGCTTTACCATTTCCA |
| AB-proAT3G62070-Fw | AGAAGTGAAGCTTGGTCTCAACCTgctctgccgtttgtcctat |
| AB-proAT3G62070-Rv | AGGGCGAGAATTCGGTCTCATGTTGAAAGAGAAGAAGAAGTAGGAA |
| CD-gAT3G62070-Fw | AGAAGTGAAGCTTGGTCTCAGGCTCCATGGCGTTCGAAGAGAGCT |
| CD-gAT3G62070-Rv | AGGGCGAGAATTCGGTCTCACTGAAGAAGCTTGACGTGTCGCA |
| B4-proAT5G11550-Fw | GGGGACAACTTTGTATAGAAAAGTTGGGTTGGAACCGTAAGGTTTA |
| B1r-proAT5G11550-Rv | GGGGACTGCTTTTTTGTACAAACTTGTcgctcactagggagactg |
| B2r-gAT5G11550-Fw | GGGGACAGCTTTCTTGTACAAAGTGGccATGATGAAAGCAAACCAGAA |
| B3-gAT5G11550-Rv | GGGGACAACTTTGTATAATAAAGTTGcTCAGGTCCCATTCTGAC |
| AB-proAT1G12845-Fw | AGAAGTGAAGCTTGGTCTCAACCTCCTCATCATCATCAAATTCTT |
| AB-proAT1G12845-Rv | AGGGCGAGAATTCGGTCTCATGTTgggtgtttctttttcttctgt |
| CD-gAT1G12845-Fw | AGAAGTGAAGCTTGGTCTCAGGCTCCATGGCTCTGTTTTCTAGC |
| CD-gAT1G12845-Rv | AGGGCGAGAATTCGGTCTCACTGAATAGTGTAGAAACCTAACAAGTA |
| B4-proAT2G41190-Fw | GGGGACAACTTTGTATAGAAAAGTTGGGcatatcgtaacaaatttgaagga |
| B1r-proAT2G41190-Rv | GGGGACTGCTTTTTTGTACAAACTTGTatttcgaacaaacccaagtaag |
| B1-gAT2G41190_ns-Fw | GGGGACAAGTTTGTACAAAAAAGCAGGCTtgATGGAGGACAAGAACAATGA |
| B2-gAT2G41190_ns-Rv | GGGGACCACTTTGTACAAGAAAGCTGGGTcCTGATAGTTCCTAATGATCTTTG |
| B4-proAT1G75900-Fw | GGGGACAACTTTGTATAGAAAAGTTGGGtctaccacgcgataggatca |
| B1r-proAT1G75900-Rv | GGGGACTGCTTTTTTGTACAAACTTGTgttgctttaatgtaatgtataaaat |
| B2r-gAT1G75900-Fw | GGGGACAGCTTTCTTGTACAAAGTGGccATGAAAGACAATTCAAGCTGG |
| B3-gAT1G75900-Rv | GGGGACAACTTTGTATAATAAAGTTGcTCAGACGAACTGATTAACAAAT |
| AB-proAT1G10850-Fw | AGAAGTGAAGCTTGGTCTCAACCTgtgtttgatagggcttctcg |
| AB-proAT1G10850-Rv | AGGGCGAGAATTCGGTCTCATGTTTGTCGATGAATGATGAATCAAT |
| CD-gAT1G10850-Fw | AGAAGTGAAGCTTGGTCTCAGGCTCCATGGCTTCTTCTTCTTCTTC |
| CD-gAT1G10850-Rv | AGGGCGAGAATTCGGTCTCACTGAAATGCTCACTTGATCATCC |
| B4-proSDD1-Fw | GGGGACAACTTTGTATAGAAAAGTTGGGAAATCATCGATGTTCTTGAAATTG |
| B1r-proSDD1-Rv | GGGGACTGCTTTTTTGTACAAACTTGTTGGAGAGAGTTAAAAAAAGGAGTT |
| B2r-gSDD1-Fw | GGGGACAGCTTTCTTGTACAAAGTGGccATGGAACCCAAACCTTTCTT |
| B3-gSDD1-Rv | GGGGACAACTTTGTATAATAAAGTTGcTCAGTTAGTCTTCAAGGTTAC |
| AB-proAT5G23400-Fw | AGAAGTGAAGCTTGGTCTCAACCTATCATCAATTCTGGCTGTCAA |
| AB-proAT5G23400-Rv | AGGGCGAGAATTCGGTCTCATGTTTTTGGGATTTGATCAGTTTAAACG |
| CD-gAT5G23400-Fw | AGAAGTGAAGCTTGGTCTCAGGCTCCATGCAAAACCTGAAATGGG |
| CD-gAT5G23400-Rv | AGGGCGAGAATTCGGTCTCACTGACTTCATTGTCTTCCTACAAGC |
| B4-proAT1G51405-Fw | GGGGACAACTTTGTATAGAAAAGTTGGGCCTTTGAAACATTAGACCTACCA |
| B1r-proAT1G51405-Rv | GGGGACTGCTTTTTTGTACAAACTTGTTACTATTCACCGCCGACTTTAA |
| B2r-gAT1G51405-Fw | GGGGACAGCTTTCTTGTACAAAGTGGccATGGAGAGACGTAACGAAG |
| B3-gAT1G51405-Rv | GGGGACAACTTTGTATAATAAAGTTGcTTACATTTTCATGATATTGGAGTT |
| AB-proAT2G39690-Fw | AGAAGTGAAGCTTGGTCTCAACCTTCTGTCCTCTGAATCCAACA |
| AB-proAT2G39690-Rv | AGGGCGAGAATTCGGTCTCATGTTGAGTTTCAGAGAAACATAAGAAG |
| AB-proAT1G71050-Fw | AGAAGTGAAGCTTGGTCTCAACCTATGGATAGGCTGAGGAAAAC |
| AB-proAT1G71050-Rv | AGGGCGAGAATTCGGTCTCATGTTGTCTATTACTCTCACCAAGAAAG |
| AB-proAT3G47675-Fw | AGAAGTGAAGCTTGGTCTCAACCTAGGATTGGGAGCTAAGGCAG |
| AB-proAT3G47675-Rv | AGGGCGAGAATTCGGTCTCATGTTACAGAGGTGCTTACTCCTTCC |
| AB-proAT2G46070-Fw | AGAAGTGAAGCTTGGTCTCAACCTAATGCAGTTGGAGAAGATGT |
| AB-proAT2G46070-Rv | AGGGCGAGAATTCGGTCTCATGTTGATGATGCAATGATCAGACC |
| AB-proAT1G76190-Fw | AGAAGTGAAGCTTGGTCTCAACCTTCTCTGGCATGAGTTTTGTG |
| AB-proAT1G76190-Rv | AGGGCGAGAATTCGGTCTCATGTTATGATTTTTGTGTGTGTGTTAC |
| AB-proAT4G38420-Fw | AGAAGTGAAGCTTGGTCTCAACCTTGTATTGTTGTCCACCTCGT |
| AB-proAT4G38420-Rv | AGGGCGAGAATTCGGTCTCATGTTACTCGCTTTGACAAGAAGAAG |
| AB-proAT4G05120-Fw | AGAAGTGAAGCTTGGTCTCAACCTgttctcatcattttgtgcaa |
| AB-proAT4G05120-Rv | AGGGCGAGAATTCGGTCTCATGTTTGAGCTCAAAAGAACATAAACC |
| CD-gAT4G05120-Fw | AGAAGTGAAGCTTGGTCTCAGGCTCCATGGCGGATAGATATGAGAAC |
| CD-gAT4G05120-Rv | AGGGCGAGAATTCGGTCTCACTGAAAAGGCATTCTTCTTACCAATAAG |
| AB-proAT3G48520-Fw | AGAAGTGAAGCTTGGTCTCAACCTTGTCAGGCTCTTGCTAG |
| AB-proAT3G48520-Rv | AGGGCGAGAATTCGGTCTCATGTTATTGTTTAATTGTTTTTTGTTCTTG |
| CD-gAT3G48520-Fw | AGAAGTGAAGCTTGGTCTCAGGCTCCATGGCATTTCTTCTGAGTTT |
| CD-gAT3G48520-Rv | AGGGCGAGAATTCGGTCTCACTGAAACGTTGTTAAGGATGTGAC |
| B4-proAT3G04840-Fw | GGGGACAACTTTGTATAGAAAAGTTGGGATGTTTCAATACGTGTTTCATCTC |
| B1r-proAT3G04840-Rv | GGGGACTGCTTTTTTGTACAAACTTGTtgtgagcttggactttgga |
| B1-gAT3G04840_ns-Fw | GGGGACAAGTTTGTACAAAAAAGCAGGCTtgATGAACACTTCGTAAGTCTC |
| B2-gAT3G04840_ns-Rv | GGGGACCACTTTGTACAAGAAAGCTGGGTcCTGAGATGTGATAGCGGAA |
| AB-proAT3G16420-Fw | AGAAGTGAAGCTTGGTCTCAACCTGTTACTCCTCAGCTTCTAGT |
| AB-proAT3G16420-Rv | AGGGCGAGAATTCGGTCTCATGTTCTTCTGCTTCTTGATAGTACTT |
| CD-gAT3G16420-Fw | AGAAGTGAAGCTTGGTCTCAGGCTCCATGGCCCAAAAGGTGG |
| CD-gAT3G16420-Rv | AGGGCGAGAATTCGGTCTCACTGAGTTGGATAAAGGACGAACATG |

| | |
|---|---|
| B4-proLTP6-Fw | GGGGACAACTTTGTATAGAAAAGTTGGGggcttcttttgaatttctttaagg |
| B1r-proLTP6-Rv | GGGGACTGCTTTTTTGTACAAACTTGTgttactttctgtttttttttttggtg |
| B2r-gLTP6-Fw | GGGGACAGCTTTCTTGTACAAAGTGGccATGAGATCTCTCTTATTAGCC |
| B3-gLTP6-Rv | GGGGACAACTTTGTATAATAAAGTTGcTTAAGATTTTCTGCTTGTCTC |
| AB-proAT5G17160-Fw | AGAAGTGAAGCTTGGTCTCAACCTAAGACTCACCATTCGTAGca |
| AB-proAT5G17160-Rv | AGGGCGAGAATTCGGTCTCATGTTttccgcgaaatcgagaga |
| CD-gAT5G17160-Fw | AGAAGTGAAGCTTGGTCTCAGGCTCCATGGATTTCCACAGCCTT |
| CD-gAT5G17160-Rv | AGGGCGAGAATTCGGTCTCACTGACTCTGCAGTCTTATTATTCCC |
| B1-proEPF2-gEPF2_ns-Fw | GGGGACAAGTTTGTACAAAAAAGCAGGCTCACTATTTGACATATTTTCTTTTG |
| B2-proEPF2-gEPF2_ns-Rv | GGGGACCACTTTGTACAAGAAAGCTGGGTAAGCTCTAGATGGCACGTGATA |
| PIP2A-Fw | CACCATGGCAAAGGATGTGGAAGCC |
| PIP2A_ns-Rv | GACGTTGGCAGCACTTCTGAA |
| **Oligos for qRT-PCR** | |
| BIN2_qPCR_Bert_Fw | GTGACTTTGGCAGTGCGAAAC |
| BIN2_qPCR_Bert_Rv | CAGCATTTTCTCCGGGAAATAATGG |
| POLAR_qPCR_FW6 | AGATGGTGGTGTCAGATG |
| POLAR_qPCR_RV6 | TTGCCTTGTCTCCAGTAG |
| POLAR_qPCR_FW7 | TATCGTGCAGTGTTATAC |
| POLAR_qPCR_RV7 | GTCATCTTCTTGTTCCTC |
| POLARLIKE1_qPCRF1 | CCTTCCGAATTGCAAAGAAGG |
| POLARLIKE1_qPCRR1 | TTGCCTCAGTTCTTCGTTTCG |
| BES1_qPCR_F | CAACCTCGCCTACCTTCAATCTC |
| BES1_qPCR_R | TTGGCTGTTCTCAAACTTAAACTCG |
| Actin2_F | GATGAGGCAGGTCCAGGAATC |
| Actin2_R | AACCCCAGCTTTTTAAGCCTTT |
| BASL_RT-PCR_Fw | CTGTCTCAGAAGAATCTGGATTC |
| BASL_RT-PCR_Rv | GAATCTACAACATTGGAACCC |
| EPF2_qPCR_Fw1 | TTGCCTCGTCCTAGTCTTC |
| EPF2_qPCR_Rv1 | CATTTCCATTTCTACCCCTCC |
| MUTE_qPCR_Fw1 | CCTAAACCGACCATCTTTCC |
| MUTE_qPCR_Rv1 | TGATCTTTACGAGCTGCCC |
| CYCA2-3_qPCR_Fw1 | GTTCCCTTGCCTCTGCTTTGC |
| CYCA2-3_qPCR_Rv1 | GCTCGCTTCTTCCTCTTGTTGG |
| CYCD7-1_qPCR_Fw1 | TCCATGCGTTTCAATGGCTAATCC |
| CYCD7-1_qPCR_Rv1 | TCCACCATCCAATTCGTCCATTCG |
| SMR4_qPCR_Fw1 | TGGAGAGGAAGACGGAGATG |
| SMR4_qPCR_Rv1 | CAAGATCTGGTGGCTGAAAG |
| DWF4_qPCR_Fw | GTGATCTCAGCCGTACATTTGGA |
| DWF4_qPCR_Rv | CACGTCGAAAAACTACCACTTCCT |

**Supplementary Data 1. Gene expression analysis of the control dataset.**

Differentially expressed genes (DEGs) for each cell cluster versus all the other clusters of the control dataset. avg_$\log_2$FC, $\log_2$ fold-change of the average expression between the given cluster and all the other clusters; pct.1, percentage of cells that express a gene within the given cluster; pct.2, percentage of cells that express a gene within all the other clusters; p_val, unadjusted *P* value from the Wilcoxon rank sum statistical test; p_val_adj, adjusted *P* value based on Bonferroni correction; pct.1-pct.2, the value of pct.1 minus pct.2. The genes were ranked by 'pct.1-pct.2' value.

**Supplementary Data 2. Differential expression pattern analysis of the stomatal development trajectory under influence of brassinolide and bikinin.**

For each gene, a negative binomial generalized additive model (NB-GAM) was fitted on the expression over pseudotime under each treatment condition, i.e., DMSO, brassinolide (BL), or bikinin (BIK). Using the conditionTest from the tradeSeq package (*29*), differential expression patterns between conditions were determined with the Wald tests. waldStat_condition2_vs_condition1, Wald test statistic of the comparison of expression pattern between condition 2 and condition 1; pvalue_condition2_vs_condition1, unadjusted *P* value from the Wald test; padj_condition2_vs_condition1, adjusted *P* value based on Benjamini & Hochberg correction ("FDR").

**Supplementary Video 1. Confocal time-lapse imaging of PL1-GFP**

Confocal time-lapse imaging of *PL1pro:gPL1-GFP/Col-0* in the abaxial epidermis of cotyledons of Arabidopsis seedlings at 2 days post germination.

**Supplementary Video 2. Confocal time-lapse imaging of BES1-GFP**

Confocal time-lapse imaging of *BES1pro:gBES1-GFP/Col-0* in the abaxial epidermis of cotyledons of Arabidopsis seedlings at 2 days post germination.
